# ER stress and mitochondrial dynamics: a tale of a wandering phosphatase DUSP28

**DOI:** 10.1101/2024.08.17.608394

**Authors:** Priyanka Bhansali, Subba Rao Gangi Setty

**Author notes:** **Corresponding author information:** Subba Rao Gangi Setty Professor Department of Microbiology and Cell Biology Indian Institute of Science CV Raman Avenue, Bangalore 560012 India /+91-9482519890.

## Abstract

Fission and fusion processes maintain the mitochondrial dynamics at a steady state and are altered during various cellular stresses. The mechanisms that regulate mitochondrial dynamics during ER stress are critically unknown. Here, we identified a dual specificity phosphatase DUSP28 with a novel role in regulating mitochondrial morphology during ER stress. Cytosolic DUSP28 translocates to mitochondria during ER stress via PERK activation and further promotes mitochondrial fission in a Drp1-dependent manner. Interestingly, the knockdown of DUSP28 activates PERK signaling and enhances ER expansion following mitochondrial elongation by enhancing S637 phosphorylation of Drp1. Overexpression of DUSP28-GFP could not rescue the loss of Drp1 function, indicating its necessity for mitochondrial fission. Further, the presence of DUSP28 on mitochondria prevents Parkin recruitment during CCCP-induced mitochondrial damage and follows STX17-dependent mitophagy. These findings illustrate the novel crosstalk between ER stress and DUSP28 phosphatase in maintaining the mitochondrial dynamics to sustain cellular stress.

**Key points:**

- PERK activation translocates cytosolic DUSP28 to mitochondria
- DUSP28 mediates Drp1-dependent mitochondrial fission
- DUSP28 phosphorylation status determines its localization and function
- DUSP28 inhibits Parkin recruitment to mitochondria upon CCCP treatment and follows STX17-mediated mitophagy

## Introduction

Intracellular signaling communicates through a series of phosphorylation and dephosphorylation of substrates performed by a set of kinases and phosphatases, respectively (Forrest et al., 2003; Manning et al., 2002; Sacco et al., 2012). Typically, phosphatases dephosphorylate either at serine/threonine or tyrosine residues on specific substrates (Alonso et al., 2004; Tonks, 2006). However, dual specificity phosphatases (DUSPs) belong to a subgroup of phospho-tyrosine phosphatases (PTPs), which can dephosphorylate both serine/threonine and tyrosine residues (Patterson et al., 2009; Pulido and Hooft van Huijsduijnen, 2008). These phosphatases share a conserved active site, HCxxxxR, with broader pocket size but shallower than the classical PTPs (Pulido and Lang, 2019). DUSPs are classified into typical and atypical based on the presence or absence of MAP kinase-interacting domain, respectively (Sacco et al., 2012). DUSP28 is an atypical phosphatase with tyrosine (Y102) instead of histidine in the active site compared to other DUSPs (Ku et al., 2017). Biochemical studies reported that DUSP28 retains very low phosphatase activity *in vitro* (Ku et al., 2017). Further, DUSP28 expression (mRNA and protein) has been shown to be elevated in hepatocellular carcinoma (HCC), pancreatic and breast cancers (Buiga et al., 2018; Lee et al., 2015; Wang et al., 2014). DUSP28 has been reported to enhance cell migration, invasion, and viability through the activation of CREB, AKT, and ERK1/2 signaling in pancreatic cancer (Lee et al., 2019). DUSP28 expression positively regulates platelet derived growth Factor A (PDGF-A) and alters the migration, invasion and proliferation of pancreatic cancer cells (Lee et al., 2017). DUSP28 is also shown to modulate the cell cycle in HCC by arresting the cells in the S phase with a concomitant decrease in G1 phase cells (Wang et al., 2014). Many of the above studies have shown that DUSP28 promotes angiogenesis and metastasis during cancer progression. However, the intracellular localization and the cellular function of DUSP28 have not been reported.

Endoplasmic reticulum (ER) stress activates unfolded protein response (UPR), a signalling cascade that monitors ER homeostasis (Hetz and Glimcher, 2009). Studies have shown that the accumulation of misfolded proteins, alteration in ER calcium levels or other cellular processes elevate ER stress (Walter and Ron, 2011). To encounter the stress, cells expand ER (Bakunts et al., 2017; Schuck et al., 2009) and activate transcriptional response through IRE1, ATF6 and PERK signaling (Walter and Ron, 2011). IRE1 and PERK are kinases that undergo trans autophosphorylation and oligomerization upon sensing ER stress. These signaling molecules activate the respective transcription factors XBP1 and ATF4, which combat the ER stress through gene transcription (Ali et al., 2011; Chen and Brandizzi, 2013; Lin et al., 2009). In contrast, ATF6 translocates to Golgi upon sensing ER stress and being processed to a transcription factor, which performs a transcriptional response (Hetz and Glimcher, 2009; Walter and Ron, 2011). Although the activation of these three arms during ER stress has been shown, the intracellular signals/conditions that trigger the arm-specific activation are unknown. Recent studies have identified several arm-specific small molecular inhibitors, such as Ceapin-A7 blocks the processivity of ATF6 to form an active transcription factor (Gallagher et al., 2016); KIRA6 inhibits the kinase activity of IRE1 by binding to its ATP binding site and prevents its phosphorylation and oligomerization (Mahameed et al., 2019); and GSK2606414 blocks the eIF2α-ATF4 signaling and alter the PERK signaling (Mahameed et al., 2019). Thus, these molecules form a potential tool to study arm-specific UPR activation.

Mitochondria regulates many cellular functions, such as migration, cell death and differentiation, immune response, and stem cell maintenance etc. (Spinelli and Haigis, 2018). During these processes, mitochondria maintain bioenergetics and respond to the cellular environment by undergoing fission and fusion events (referred to here as mitochondrial dynamics) or clearance through mitophagy (Giacomello et al., 2020). Several known proteins regulate the mitochondrial fission (Fis1, MiD49, Mff and Drp1) and fusion (Mfn1, Mfn2 and Opa1), which undergo a variety of post-translational modifications corresponding to the cellular status (Giacomello et al., 2020). Studies have shown that ER marks the fission site on mitochondria to which Drp1 will be recruited by mitochondrial adapters such as Fis1, MiD49/MiD51 and Mff (Friedman and Voeltz, 2011; Loson et al., 2013). Further, oligomerization of Drp1 causes mitochondrial fission at the ER-mitochondrial contact site (Frohlich et al., 2013). Interestingly, Drp1 fission activity is regulated by its phosphorylation at Serine (S) 616 and S637, in which p-S616 promotes fission and pS637 inhibits fission (van der Bliek et al., 2013). Several kinases like protein kinase C isoform delta (PKCδ) (Zaja et al., 2014), Rho-associated protein kinase (ROCK) (Brand et al., 2018), Cdk1/Cdk5, CaM kinase Iα (Han et al., 2008) or PINK1 (Han et al., 2020) phosphorylates Drp1 at S616 residue, whereas p-S637 mediated by protein kinase A (PKA) (Chang and Blackstone, 2007). In contrast, the dephosphorylation of Drp1 at p-S637 residue is possibly regulated by calcineurin (Cereghetti et al., 2008; Frohlich et al., 2013). On the other hand, mitophagy helps in clearing damaged mitochondria, which occurs majorly through the PINK1/Parkin-mediated pathway (McQuiston and Diehl, 2017).

ER and mitochondria are known to crosstalk and associate through their mitochondrial-associated membrane (MAM) contact sites (Rowland and Voeltz, 2012). PERK has been reported to be present at MAMs and facilitate inter-organelle communication, including their dynamics and homeostasis (Almeida et al., 2022; Lebeau et al., 2018; Munoz et al., 2013; Verfaillie et al., 2012). These organelles vary their morphology and activate signaling during stress conditions to adopt or maintain cellular homeostasis. However, the regulation of mitochondrial dynamics during ER stress is critically unknown. Interestingly, the PERK arm promotes the adaptive remodeling of mitochondrial membranes through phosphatidic acid and enhances its elongation during ER stress (Perea et al., 2023). Additionally, ER signaling/contacts are known to alter the mitochondrial dynamics by modulating Drp1 activity. For example, calcineurin, a Ca^2+^-dependent serine-threonine phosphatase, associates with the PERK signaling and regulates mitochondrial dynamics by dephosphorylating Drp1 (Cereghetti et al., 2008; Cribbs and Strack, 2007). Phosphoglycerate mutase family member 5 (PGAM5) phosphatase localizes to the ER-mitochondria interface and regulates mitochondrial dynamics during necrosis (Peng et al., 2022; Wang et al., 2012) or through MFN2 (Nag et al., 2023). However, the molecular mechanism linking ER stress and mitochondria dynamics remains unknown.

In this study, we identified an atypical phosphatase DUSP28 that senses ER stress and regulates mitochondrial dynamics through Drp1. The function of DUSP28 is possibly controlled through its phosphorylation, which inhibits its translocation to mitochondria. Upon PERK activation, DUSP28 localizes to mitochondria and mediates Drp1-dependent fission. Depletion of DUSP28 activates PERK and results in mitochondrial elongation with a concomitant increase in S637 phosphorylation of Drp1. Expression of Drp1(S637D) mutant in siDUSP28 cells restored the mitochondrial fission. However, overexpression of DUSP28 in Drp1 knockdown cells or depletion of Drp1 recruiters (Fis1, MiD49 or MiD51) did not rescue the defective mitochondrial fission in HeLa cells. Further, DUSP28 localization to mitochondria prevents Parkin recruitment and facilitates STX17-mediated mitophagy during CCCP (carbonyl cyanide mDchlorophenylhydrazone)-induced mitochondrial damage. In a nutshell, these studies link DUSP28 to ER stress and regulate mitochondrial dynamics through molecular signaling.

## Results

### Ectopic expression of DUSP28 localizes to mitochondria and causes its fragmentation

To examine the intracellular function of DUSP28, we studied its localization as a C-terminal GFP-fusion (**Fig. 1A**) due to the lack of commercial antibodies suitable for immunofluorescence microscopy (IFM). Transient expression of DUSP28-GFP in HeLa cells showed three different patterns of localization, which includes cytosolic (C, 56±3% cells), membranous and cytosolic (C+M, 26±6% cells), and membranous only (M, 16±3% cells) within the cells (**Fig. 1B**). Investigation to identify the nature of membranous location of DUSP28-GFP with organelle specific markers showed its complete colocalization with mitochondrial marker, TOM20 (Pearson’s coefficient, *r*=0.59±0.09) (**Fig. 1B**). Further, mitochondrially localized DUSP28-GFP showed partial colocalization with multiple organelles such as with ER, early endosomes, and lysosomes (marked with KDEL-RFP, EEA1 and LAMP1, respectively) but no apparent colocalization with *trans* Golgi (labelled with p230) (*r -* values of membranous DUSP28-GFP with KDEL-RFP=not able to be determined; EEA1= 0.18±0.06; and LAMP1= 0.21±0.05) (**Fig. 1C**). The live-cell imaging of cells expressing DUSP28-GFP showed constant colocalization with the mitochondrial marker DsRed2-Mito-7 (referred to here as DsRed-Mito) throughout the duration of time lapse imaging (**Fig. S1A** and **Video S1**), suggesting that DUSP28-GFP localizes primarily to mitochondrial membranes in addition to cytosol upon overexpression. Analysis of the DUSP28 cDNA sequence for the mitochondrial targeting motifs identified a Kozak sequence internally starting from the 87^th^ amino acid (aa) but no putative mitochondrial targeting motifs. We predicted that this internal start site may produce a shorter isoform of DUSP28 (referred to here as sDUSP28, having 87-176 aa of wild type) (**Fig. 1A**). Immunoblotting analysis of cells expressing DUSP28-GFP showed the presence of sDUSP28-GFP form, which has equivalent size to the protein expressed by sDUSP28-GFP plasmid in HeLa cells (**Fig. S1B**). Interestingly, expression of sDUSP28-GFP mostly showed its localization to mitochondria (labelled with DsRed2-Mito-7) in HeLa cells (**Fig. 1B**). Consistently, the population of cells with mitochondrial sDUSP28-GFP (70±4%) was enhanced compared to the full length DUSP28-GFP (44±3%) (**Fig. 1D**). Additionally, increasing expression of DUSP28-GFP showed enhanced mitochondrial localization ratiometrically (**Fig. S1C**), indicating that the expression levels of DUSP28-GFP also regulate its localization to the mitochondria.

**Figure 1.**
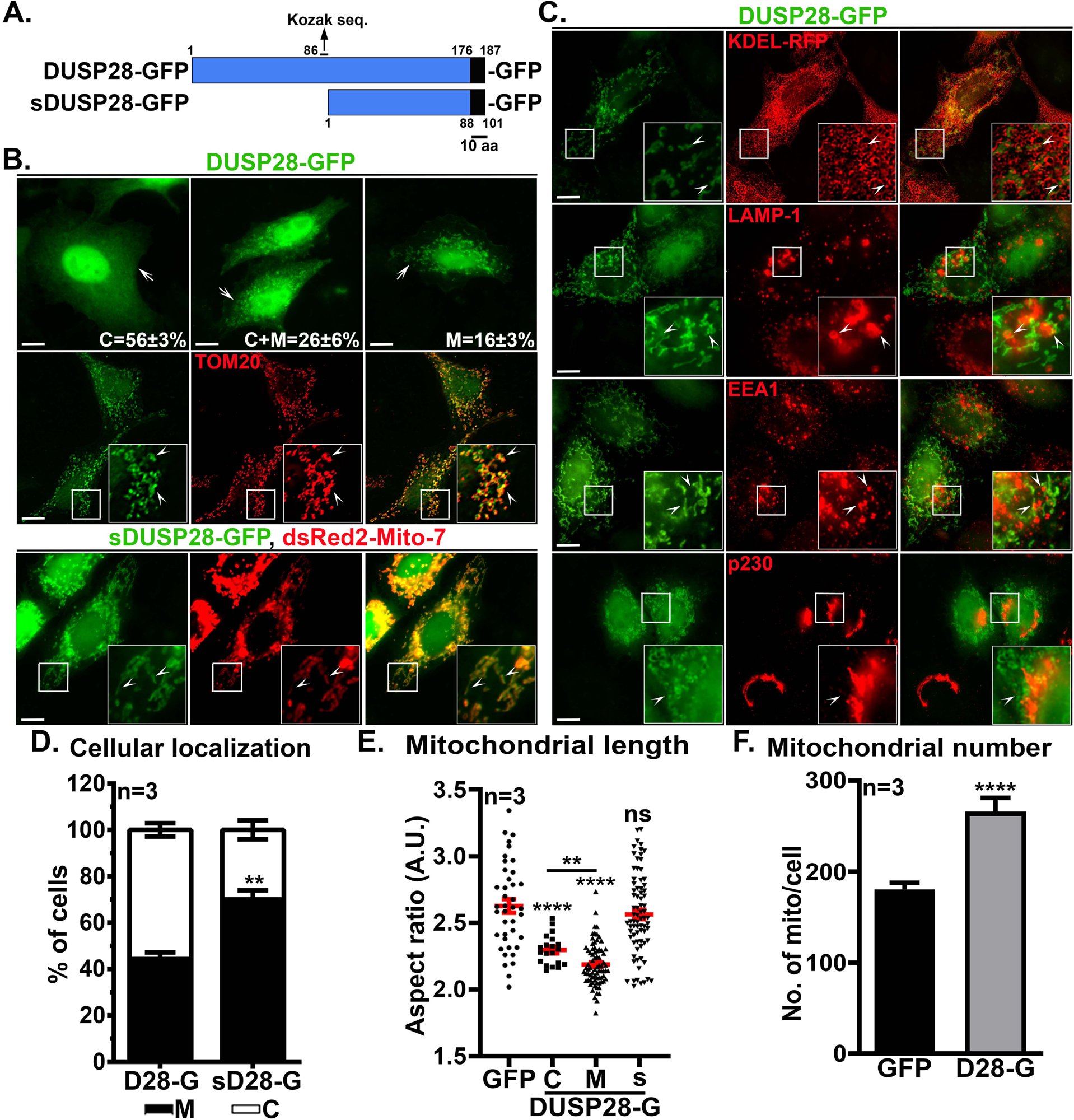
Transiently overexpressed DUSP28-GFP localizes to mitochondria and alters its morphology. **(A)** Schematic representation of full length and shorter isoform (s) of C-terminal GFP tagged DUSP28. Scale, 10 aa. **(B)** Localization of DUSP28-GFP in HeLa cells showed three different patterns (arrows): cytosol (C) alone (major population), both cytosol and membranous localization (C+M) and membranous (M) alone. The membranous DUSP28 colocalizes with TOM20, a mitochondrial outer membrane receptor (arrowheads). sDUSP28-GFP expression in HeLa cells showed a major localization to the mitochondrial membranes and colocalized with the dsRed2-Mito-7 marker (arrowheads). **(C)** Analysis of membranous DUSP28-GFP localization in HeLa cells by co-staining with different organelle specific markers: KDEL-RFP for endoplasmic reticulum, LAMP-1 for lysosomes, EEA1 for early endosomes and p230 for *trans*-Golgi. Arrowheads point to the localization of mitochondrial DUSP28-GFP with respect to other organelles. All insets of IFM images are amplified white box areas. Scale 10 μm. **(D-F)** Graphs representing the quantification of different parameters measured in DUSP28-GFP expressing HeLa cells. The percentage of cells showing the cytosolic (C) or mitochondrial (M) localization of DUSP28-GFP **(D).** The length of mitochondria was measured using aspect ratio (arbitrary units, A.U.) in cells expressing GFP, cytosolic/membrane bound DUSP28-GFP or membrane bound sDUSP28-GFP **(E).** The number of mitochondria in cells expressing GFP or DUSP28-GFP **(F).** n=3. All values are presented as mean±s.e.m. **p≤0.01, ****p≤0.0001 and ns= not significant.

Visual observations of IFM images directed our attention towards a change in mitochondrial morphology when DUSP28-GFP localizes to mitochondria compared to the cytosol (**Fig. 1B**, **1C** and **S1D**). Quantification of mitochondrial morphology using aspect ratio (AR, a ratio of the major axis to minor axis) showed a significantly decreased value in cells expressing DUSP28-GFP compared to GFP alone, indicating shorter mitochondria (**Fig. 1E**). The AR of mitochondria was further decreased in cells expressing mitochondrial localized DUSP28 compared to its cytosolic form (**Fig. 1E**). The reduced AR values of mitochondria suggest the enhanced mitochondrial fragmentation in the cells expressing DUSP28-GFP. However, the expression of sDUSP-28 showed no major change in mitochondrial AR values compared to GFP expressing cells (**Fig. 1E**), suggesting that N-terminal (1-87 a.a) do not regulate the targeting of DUSP28 to mitochondria but may require for its function (see below). In line with these results, the cells expressing DUSP28-GFP showed enhanced mitochondrial number compared to GFP expressing cells (**Fig. 1F**). IFM analysis of mitochondrial DUSP28-GFP expressing cells showed a slight expansion of tubular ER (% tubular ER = 59.16±6.03 in DUSP28-GFP expressing cells and 52.03±6.31 in non-transfected HeLa cells), a noticeable aggregation of LAMP1 and EEA1-positive compartments and no apparent changes in the morphology of Golgi compared to the cytosolic DUS28-GFP expressing cells (**Fig. 1C** and **S1D**). These studies indicate that DUSP28-GFP localizes to mitochondria and alters its morphology by fragmentation.

DNA sequencing followed by amino acid (aa) analysis of DUSP28 amplified from HeLa cell lysate showed two aa changes, S52P and C99R, compared to the reported sequence (NP_001357394.1) in the NCBI database (**Fig. S1E**). Additionally, these mutations were consistently present in the DUSP28 sequence amplified from other cell types, such as HEK 293T cells. To be assertive that the mitochondrial fragmentation observed with mitochondrial DUSP28-GFP was not due to these mutations, we restored the mutations individually and in combination, and measured their localization to mitochondria in the HeLa cells. Quantification of cells expressing mutant or wild type form of DUSP28-GFP showed the same proportion of cells with mitochondrial localization (**Fig. S1F**), suggesting that these mutants localize similarly to DUSP28^WT^-GFP (**Fig. S1F**). Overall, these intra-cellular localization studies suggest that DUSP28-GFP majorly exists in cytosol form, with a fraction of cells displaying mitochondrial localization with fragmentation.

### ER stress drives the translocation of DUSP28 to mitochondria

The transient transfection of DUSP28-GFP in HeLa showed a variable expression that alters the population of cells showing mitochondrial localization. To reduce this variability, we prepared HeLa cells stably expressing DUSP28-GFP using lentiviral transduction (referred to here as HeLa:DUSP28-GFP). Surprisingly, 100% of the cell population showed cytosolic localization of DUSP28-GFP post-puromycin selection. Further, the majority of the cells displayed few DUSP28-GFP puncta in the cytosol (**Fig. 2A**) (discussed later). IFM characterization of HeLa:DUSP28-GFP stable cells showed no drastic difference in the organization and morphology of different cellular organelles (**Fig. S2A**). Various endogenous protein levels of different organelles are not altered in HeLa:DUSP28-GFP stables compared to its parental cell line (**Fig 2B**). Nevertheless, the stable cells displayed reduced mitochondrial AR values compared to plain HeLa cells (**Fig. S2B**). The slightly enhanced mitochondrial fragmentation in the stable cells could be due to the presence of a small cohort of DUSP28-GFP on mitochondria as observed by biochemical fractionation (see **Fig. S2F**), which may be masked by the cytosolic signal in the fluorescence microscopy images (**Fig. S2A**).

**Figure 2.**
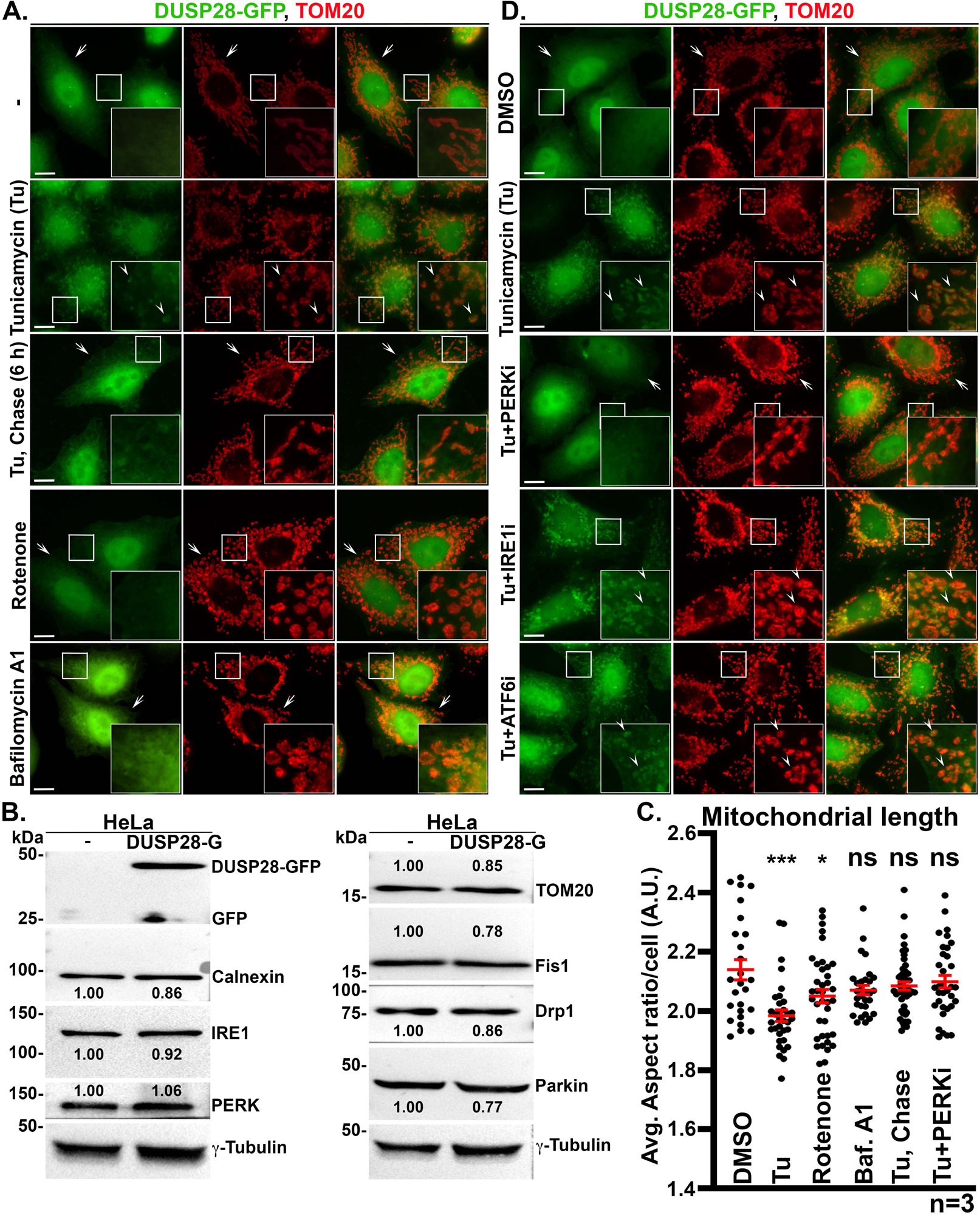
DUSP28 translocates to mitochondria upon sensing ER stress. **(A)** Stably expressed DUSP28-GFP localizes to the cytosol in HeLa cells. Treatment of cells with various stress inducing agents such as tunicamycin (Tu, 10 μm for 6 h) for ER stress, rotenone (500 nM for 6 h) for mitochondrial stress and bafilomycin A1 (5 mM for 6 h) for lysosomal stress (by altering the acidity), showed the translocation of DUSP28-GFP to mitochondria and colocalized with TOM20 only in Tu treated cells. DUSP28-GFP localization to the cytosol was reversed upon replenishing Tu with normal media for 6 h. Arrows indicate the cytosolic DUSP28-GFP, and arrowheads point to the mitochondrial DUSP28-GFP. Scale 10 μm. **(B)** Immunoblotting analysis of stably expressing DUSP28-GFP cells and its parental HeLa cells. The endogenous expression of several ER and mitochondrial proteins is unaffected in the stable cells. γ-tubulin is used as a loading control. The fold change in expression was indicated on the blots. **(C)** Plot representing the quantification of mitochondrial length (aspect ratio in arbitrary units, A.U.) in cells treated with various drugs as indicated. All values are presented as mean±s.e.m. n=3. *p≤0.05, ***p≤0.001, and ns=not significant. **(D)** IFM images of stably expressing DUSP28-GFP HeLa cells treated with DMSO, Tu, Tu+PERKi, Tu+IRE1i or Tu+ATF6i, and co-stained with TOM20 (ATF6i - 5 μM; IRE1i - 1 μM; PERKi - 100 nM for 6 h). Arrows indicate the cytosolic DUSP28-GFP, and arrowheads point to the mitochondrial DUSP28-GFP. All insets of IFM images are amplified white box areas. Scale 10 μm.

To investigate the cellular condition to which cytosolic DUSP28-GFP drives to the mitochondria, we treated the stable cells with small molecules that induce a variety of organelle stress such as ER, mitochondria or lysosomal stress. Treatment of stable cells with a mitochondrial stress inducer, rotenone (500 nM for 6 h) or lysosomal stress through bafilomycin A1 (5 mM for 6 h) showed no change in the translocation of DUSP28-GFP from the cytosol to mitochondria (**Fig. 2A**). Note that the altered mitochondrial morphology observed in the drug treated cells (**Fig. 2A**) is indicative of action of these compounds on mitochondria. The treatment of stable cells with an ER stress inducer tunicamycin (10 μM for 6 h) displayed the translocation of DUSP28-GFP to mitochondria (**Fig. 2A**). To test the specificity of ER stress in translocating DUSP28-GFP to mitochondria, we treated the DUSP28-GFP expressing HeLa cells with other ER stress inducers such as thapsigargin (500 nM for 6 h) or DTT (1 mM for 6 h). The population of mitochondrial localized DUSP28-GFP cells were enhanced with all the ER stress inducers, suggesting a role for general ER stress in driving DUSP28-GFP to mitochondria (**Fig. S2C**). To test whether the translocation of DUSP28-GFP from the cytosol to mitochondria is reversible, we removed tunicamycin and chased the cells for 6 h in plain medium. Interestingly, upon removal of the ER stress inducer, DUSP28-GFP restored its localization to the cytosol, similar to the DMSO treatment (**Fig. S2C**). To test the possibility that the DUSP28 levels may contribute to the ER stress dependent translocation to mitochondria, we analyzed the protein levels by IFM and immunoblotting. Interestingly, none of these drug treatments altered the levels of stably expressing DUSP28-GFP in HeLa cells (measured by CTCF, corrected total cell fluorescence) or its protein levels (**Figs. S2D** and **S2E**). Further, the treatment of stable cells with an inhibitor of proteasome degradation, MG132 (5 μM for 6 h), enlightened the presence of sDUSP28-GFP form (**Fig. S2E**), suggesting that the cells express sDUSP28-GFP, which is prone for proteasomal degradation. Isolation of crude mitochondria using subcellular fractionation (Wieckowski et al., 2009) showed enrichment of DUSP28-GFP to the mitochondrial membrane fraction upon treatment with tunicamycin compared to DMSO treatment (**Fig. S2F**). Altogether, these data demonstrate that the translocation of DUSP28-GFP from cytosol to mitochondria is dependent on ER stress.

Next, we investigated if the stable expression of DUSP28-GFP and its translocation to mitochondria would affect the mitochondrial morphology and dynamics. Treatment of stable cells with tunicamycin translocated the DUSP28-GFP to mitochondria and resulted in lower AR value compared to its cytosolic form of DUSP28-GFP (**Fig. 2C**). Upon rescuing the cells from tunicamycin-induced ER stress, the DUSP28-GFP relocated to the cytosol and restored the AR value of mitochondria as observed in normal cells (**Figs. 2A** and **2C**). Interestingly, a slight reduction in the AR value of mitochondria upon treatment with rotenone compared to control cells, but no translocation of DUSP28-GFP to mitochondria indicates the presence of other mitochondrial fission mechanisms (**Figs. 2A** and **2C**). These studies suggest that DUSP28-GFP migrates to mitochondria only under the ER stress conditions and enhances its fragmentation.

ER stress activates the UPR pathway, which consists of ATF6, IRE1 and PERK arms (Walter and Ron, 2011). To test if the DUSP28-GFP translocation to mitochondria is mediated by general ER stress or specific to single-arm activation, we employed small molecular inhibitors against each pathway and subjected the DUSP28-GFP stable cells to tunicamycin (for 6 h). Treatment of the stable cells with Ceapin-A7 (inhibitor of ATF6 pathway, ATF6i) (Gallagher et al., 2016), KIRA6 (inhibitor of IRE1 pathway, IREi) or GSK2606414 (inhibitor of PERK pathway, PERKi) (Mahameed et al., 2019) inhibited the translocation of DUSP28-GFP to mitochondria with PERKi but not with ATF6i or IRE1i (**Fig. 2D**). The specific effect of the PERKi was further confirmed by using immunoblotting analysis of cell lysates (**Fig. S2G**). Additionally, we did not observe any increase in DUSP28 transcript levels during tunicamycin induced ER stress (**Fig. S2H**). Thus, these studies suggest that the translocation of DUSP28-GFP to mitochondria is dependent on PERK activation.

### The active site of DUSP28 plays a regulatory role for its mitochondrial localization

DUSP family members share homology in the catalytic domain and possess the HCxxGxxR sequence as their signature motif (x - any aa residue) (Alonso and Pulido, 2016; Pulido and Lang, 2019). However, DUSP28 contains a YCKNGRSR motif, an atypical active site with unusual bulky residues that contributes to steric hindrance and is thus reported to have exceptionally low phosphatase activity *in vitro* (Ku et al., 2017). Another unique feature of DUSP28’s active site is the presence of tyrosine (Y) as the first residue compared to histidine (H), found in all the other DUSP family members (Ku et al., 2017). Here, we tested the importance of tyrosine in the active site of DUSP28 in regulating its mitochondrial localization and/or fragmentation by performing site-directed mutagenesis. To begin with, we first mutated the tyrosine to histidine (Y102H) (**Fig. 3A**) to mimic the active site of other DUSPs. Surprisingly, the DUSP28^Y102H^-GFP mutant was insensitive to tunicamycin induced mitochondrial translocation and localized majorly to cytosol compared to DUSP28^WT^-GFP (**Figs. 3B** and **3D**). Further, the DUSP28^Y102H^-GFP mutant expressing HeLa cells did not display mitochondrial fragmentation (**Fig. 3B**), indicating the inactive nature of the DUSP28^Y102H^ mutant as reported previously (Ku et al., 2017).

**Figure 3.**
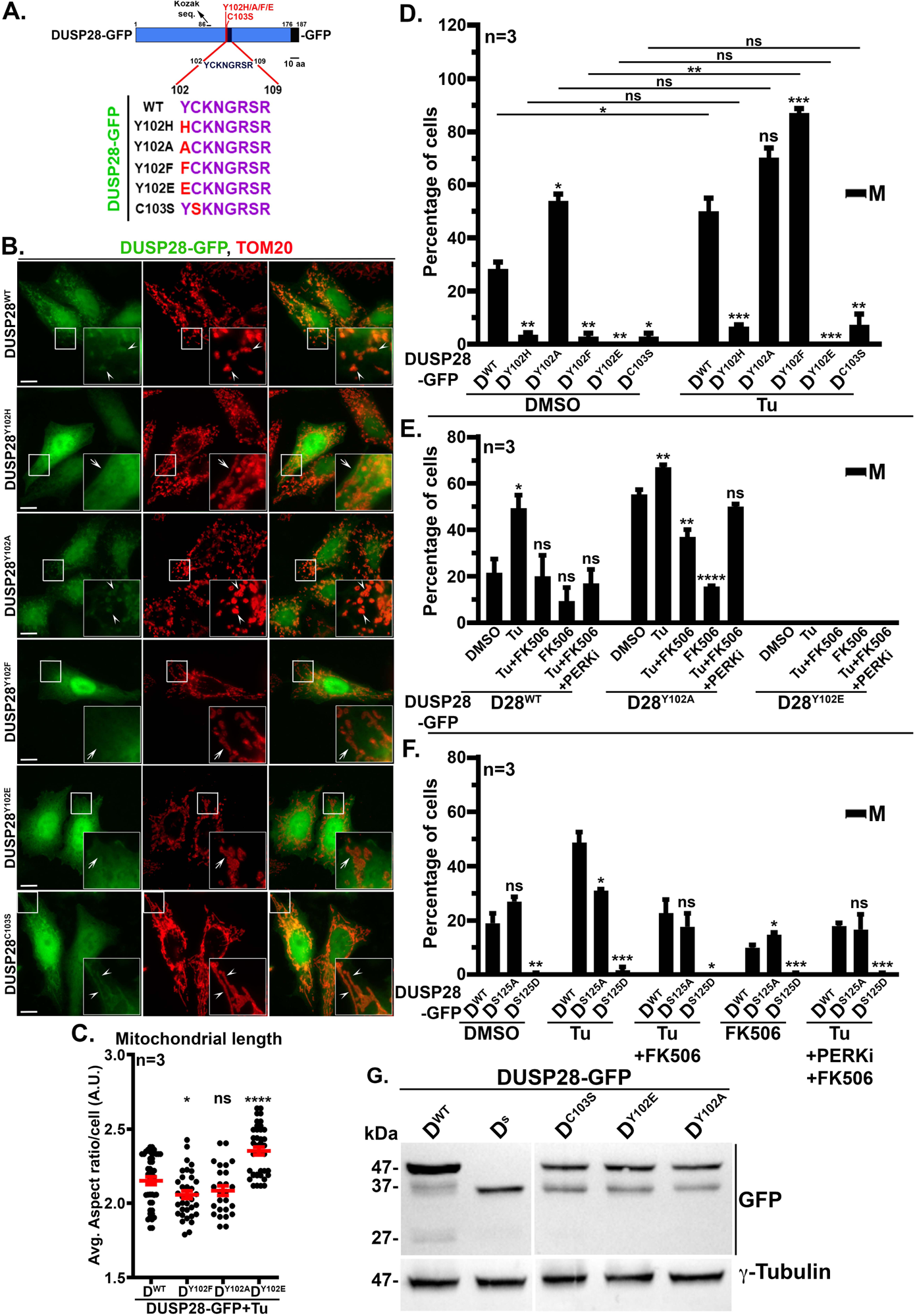
The putative active site of DUSP28 regulates its localization to the mitochondria. **(A)** Schematic diagram of DUSP28 structure illustrating its active site residues (^102^YCKNGRSR^109^), which was mutated (red colour) as indicated. **(B)** IFM images of HeLa cells expressing DUSP28^WT^-GFP or its mutants DUSP28^Y102H^-GFP, DUSP28^Y102A^-GFP, DUSP28^Y102F^-GFP, DUSP28^Y102E^-GFP, DUSP28^C103S^-GFP individually and co-stained for TOM20. All images were acquired post 24[h of transfection. Arrowheads point to the mitochondrial DUSP28-GFP, and arrows indicate the cytosolic DUSP28-GFP. All insets are amplified white box areas. Scale 10 μm. **(C)** Quantification of mitochondrial length (aspect ratio in arbitrary units, A.U.) in cells expressing DUSP28-GFP or its mutants post tunicamycin treatment (Tu, 10 μm for 6 h). n=3. **(D-F)** Percentage of cells with mitochondrial localization of DUSP28^WT^-GFP and its mutants as indicated in the presence and absence of tunicamycin (Tu, 10 μm for 6 h) **(D)**. Percentage of cells with mitochondrial localization of DUSP28-GFP WT, Y102A or Y102E mutants under various treatment conditions as mentioned (see materials and methods) **(E)**. Percentage of cells with mitochondrial localization of DUSP28-GFP WT, S125A or S125D mutants with various treatment conditions as mentioned (see materials and methods) **(F)**. All values are presented as mean±s.e.m. n=3. *p<0.05, **p≤0.01, ***p≤0.001, ****p≤0.0001 and ns=not significant. **(G)** Immunoblotting analysis of HeLa cell lysate expressing DUSP28-GFP WT or its individual mutants, or sDUSP28-GFP to verify the protein stability. The blot was probed with an anti-GFP antibody. Note that the expression of sDUSP28-GFP was observed in all mutants expressing DUSP28-GFP cells. GFP cleavage was observed in cell lysate expressing DUSP28^WT^-GFP. γ-tubulin was used as the loading control.

To study the effects of altering the pocket size residues in the active site of DUSP28, we replaced tyrosine with a smaller-sized amino acid alanine (Y102A) (**Fig. 3A**). The expression of DUSP28^Y102A^-GFP mutant showed an enhanced percentage of HeLa cells displaying mitochondrial DUSP28, which was further increased with the treatment of tunicamycin (**Figs. 3B** and **3D**). Further, DUSP28^Y102A^-GFP mutant expression caused mitochondrial fragmentation compared to DUSP28^WT^ (**Figs. 3B, 3C** and **3D**), indicating the enhanced activity of DUSP28 upon changing of tyrosine to alanine in the active site. An additional possibility is that Y102 of DUSP28 might be regulated through phosphorylation, due to which DUSP28^Y102A^-GFP mutant mimics a phospho-dead form, showing an increased mitochondrial localization and fragmentation. To test this hypothesis, we generated another phospho-dead mutant of DUSP28 by replacing tyrosine with phenylalanine (Y102F) (**Fig. 3A**). Astonishingly, DUSP28^Y102F^-GFP mutant expression in HeLa cells showed reduced localization to mitochondria compared to DUSP28^WT^; however, the mutant localized to mitochondria upon induction of ER stress and drastically increased their cell population much higher than that of DUSP28^WT^ (**Figs. 3B, 3C** and **3D**). Additionally, the DUSP28^Y102F^-GFP mutant expression caused the mitochondrial fragmentation similar to DUSP28^Y102A^-GFP mutant in HeLa cells (**Fig. 3C**). In line with these results, the phospho-mimetic mutant of DUSP28^Y102E^-GFP (**Fig. 3A**) altogether lost its mitochondrial localization irrespective of induced ER stress (**Fig. 3D**). Thus, DUSP28^Y102E^-GFP mutant expression showed its cytosolic localization in 100% cell population with reduced mitochondrial fragmentation compared to DUSP28^WT^-GFP cells (**Figs. 3B, 3C** and **3D**). These results suggest a possible post-translation modification (preferably phosphorylation) at Y102 of DUSP28, which might be controlling the activity of DUSP28, including its localization. This active site-specific modification of DUSP28 phosphatase needs to be investigated in future.

It has been reported that mutation at C103S in the active site of DUSP28 results in catalytically inactive in an *in vitro* phosphatase activity assay (Ku et al., 2017). DUSP28^C103S^-GFP mutant in HeLa cells was localized majorly to the cytosol and showed insensitive to the ER stress induction for its translocation to the mitochondria (**Figs. 3B** and **3D**). We observed the mitochondrial localization of DUSP28^C103S^-GFP in approximately 2% of cells with no apparent change in mitochondrial fragmentation (**Fig. 3B** and **3D**). Unexpectedly, these cells visually showed an apparent change in mitochondrial morphology (**Fig. 3B**). Next, we tested the stability of DUSP28 mutants’ expression in HeLa cells, which was not altered majorly, indicating their stability similar to DUSP28^WT^ (**Fig. 3G**). Overall, the active site of DUSP28 regulates not only mitochondrial fragmentation but also its translocation to the mitochondria, possibly through tyrosine phosphorylation.

Our studies showed that the PERK arm regulates the localization of DUSP28 and its dependent role in mitochondrial fragmentation. Earlier reports indicated that the PERK arm also activates calcineurin phosphatase (Wang et al, 2013). Here, we tested the role of calcineurin in regulating DUSP28 function. Cells expressing DUSP28-GFP were treated with a calcineurin inhibitor FK506 (Dumont, 2000), which drastically affected the localization of DUSP28 both at the basal and ER stress induced conditions (**Fig. 3E**). Additionally, no additive effect was observed when the cells treated with in combination of drugs, FK506 and PERKi, as PERK regulates calcineurin (**Fig. 3E**). Moreover, the trend was replicated for DUSP28^Y102A^-GFP, which normally has more cells with mitochondrial localization when compared to the DUSP28^WT^-GFP under basal conditions. Additionally, DUSP28^Y102E^-GFP showed no change in its cytosolic localization in HeLa cells (**Fig. 3E**). These studies indicate a role for calcineurin in modulating the activity of DUSP28. Calcineurin especially dephosphorylates the phospho-serine/threonine residues, which are not present in the active site of DUSP28. Here, we hypothesized that calcineurin possibly dephosphorylates DUSP28 at a site other than the active site to regulate its mitochondrial localization.

Next, we scanned the DUSP28 protein sequence for serine or threonine residues that may be potential sites for calcineurin phosphatase activity. The bioinformatics analysis using NetPhos-3.1 software (https://services.healthtech.dtu.dk/service.php?NetPhos-3.1) found Serine at the 125^th^ position in the DUSP28 sequence, which potentially be phosphorylated. We generated DUSP28^S125A^ and DUSP28^S125D^ mutants correspondingly to the phospho-dead and phospho-mimetic nature of S^125^ residue, respectively. We expressed these mutants in HeLa cells and scored their mitochondrial localization in the presence and absence of tunicamycin and FK506 (**Fig. 3F**). These results demonstrated that phospho-mimetic mutant of DUSP28 was unable to localize to mitochondria irrespective of the cellular status, suggesting S125 residue in DUSP28 is possibly dephosphorylated in calcineurin-dependent manner (**Fig. 3F**). Unexpectedly, the phospho-dead mutant of DUSP28^S125A^ did not behave as predicted i.e., the maximum population of cells should show mitochondrial localized DUSP28-GFP (**Fig. 3F**). However, the mutant DUSP^S125A^ localized as similar to DUSP28^WT^. The only plausible explanation owing to these results is that S125 in DUSP28 is regulated by calcineurin; however, this site is not solely responsible for regulating DUSP28 localization to mitochondria. Further, dephosphorylation by calcineurin accounts for the first step, which is further dependent on the active site of DUSP28(Y102E) that does not respond to any stimulation. Hence, the activity of DUSP28 phosphatase is regulated by phosphorylation at multiple sites.

### DUSP28 depletion in HeLa cells alters the mitochondrial dynamics

Overexpression of DUSP28 enhances its residence on mitochondria and increases mitochondrial fragmentation. To test the role of endogenous DUSP28, we depleted the HeLa cells with gene-specific siRNA and evaluated the changes in organellar distribution/morphology. The knockdown of DUSP28 in HeLa cells reduced the transcript levels approximately by 70% (**Figs. S3A** and **S3B**) and caused the elongation of mitochondria (**Fig. 4A**). Consistently, the aspect ratio and average area of mitochondria per cell were significantly increased in the DUSP28 knockdown compared to control siRNA treated cells (**Fig. 4B**). Further, live imaging microscopy experiments showed an increase in mitochondrial length, area, and aspect ratio in DUSP28 depleted compared to control cells (**Figs. S3C-S3F, and Video S2 and S3**). These results demonstrate a role for DUSP28 in regulating mitochondrial dynamics at the basal level. To negate the observed alterations in mitochondrial morphology of DUSP28 knockdown cells is not due to the changes in mitochondrial membrane potential, we employed 10-N-Nonyl acridine orange (NAO), Tetramethylrhodamine, ethyl ester (TMRE) and MitoTracker™ Red CMXROS, which measures the mitochondrial mass and membrane potential respectively. The DUSP28 knockdown cells showed no significant changes in the fluorescent intensity of these probes compared to control cells (**Figs. S3G-S3I**).

**Figure 4.**
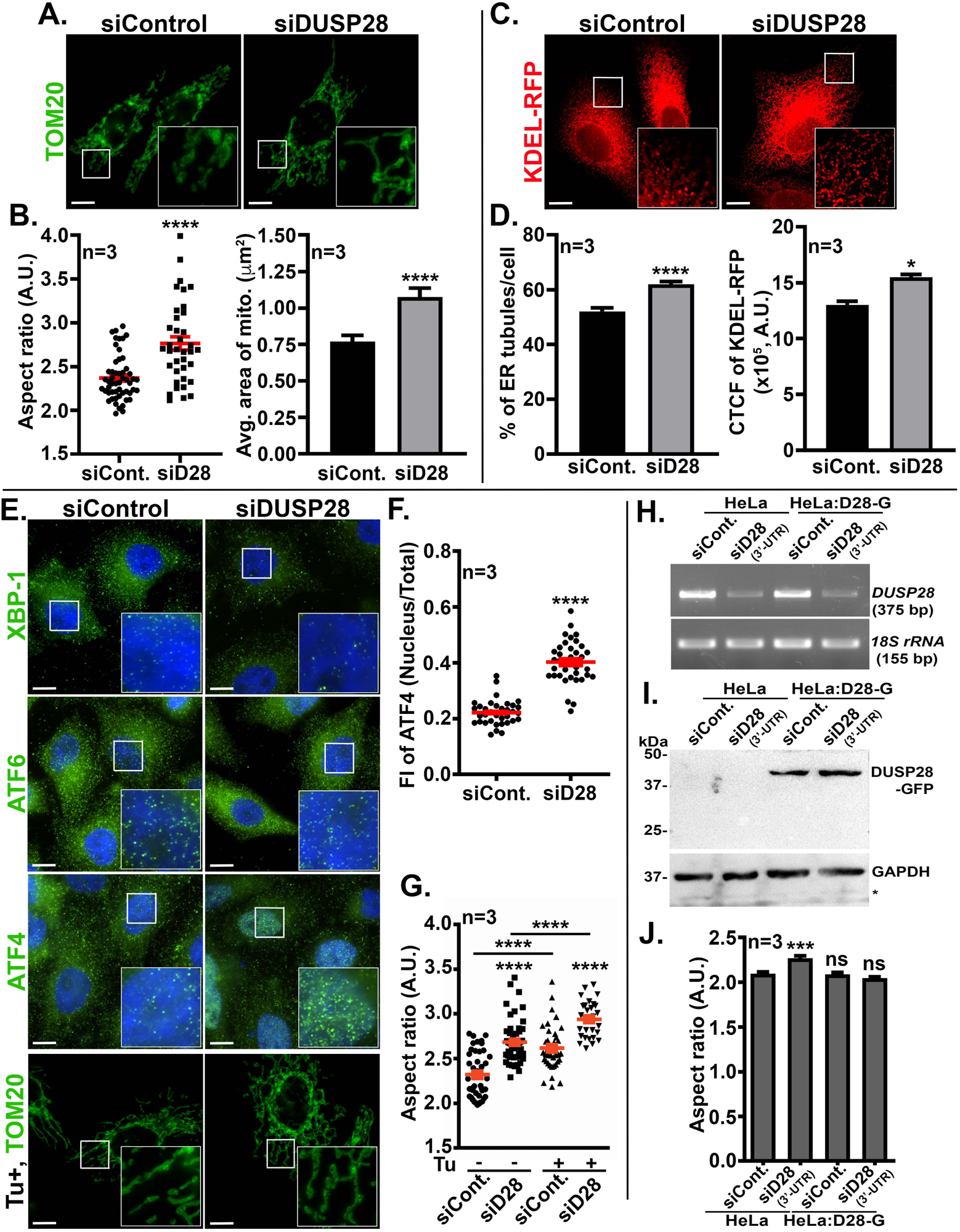
Depletion of DUSP28 in HeLa cells perturbs the mitochondrial dynamics and causes ER stress. **(A)** IFM images of siControl and siDUSP28 transfected cells that were immunostained for mitochondrial protein TOM20. **(B)** The images of **A** were quantified to measure the average aspect ratio (arbitrary units, A.U.) and the area of mitochondria per cell, presented as separate graphs. n=3. **(C)** IFM analysis of control and DUSP28 knockdown cells that were transfected with KDEL-RFP to mark the ER. The images were quantified and measured the percentage area of ER tubules per cell and the fluorescence intensity (CTCF) of KDEL-RFP per cell, plotted and shown in **D.** n=3. **(E)** IFM images of control and DUSP28 depleted cells that were stained for ER stress transcription factors, XBP-1, ATF6 and ATF4 to represent the activation of IRE1, ATF6 and PERK pathways, respectively. **(F)** Graph representing the nuclear localization of ATF4 fluorescence intensity (FI) in the nucleus vs total in siControl and siDUSP28 transfected cells. n=3. **(G)** Graph representing the quantification of mitochondrial aspect ratio (arbitrary units, A.U.) per cell in control and DUSP28 knockdown cells in the absence and presence of tunicamycin (Tu). n=3. Representative IFM images of control and DUSP28 knockdown cells stained for TOM20 are shown in the bottom most panel of **E**. All insets of IFM images are amplified white box areas. Scale 10 μm. **(H-J)** Rescue experiments to demonstrate the specific effect of DUSP28 knockdown on mitochondria using 3’-UTR specific siRNA. The siRNAs were transfected both in HeLa and DUSP28-GFP stable HeLa cells to knock down the endogenous DUSP28. The mRNA levels of DUSP28 were analyzed by agarose gel electrophoresis, and *18S rRNA* was used as the internal control **(H)**. **(I)** Immunoblotting analysis of DUSP28-GFP levels in control and DUSP28-knockdown (3’-UTR) cells. GAPDH is used as a loading control. *, indicates the non-specific band recognized by anti-GAPDH antibody. **(J)** Graph representing the quantification of mitochondrial aspect ratio (arbitrary units, A.U.) in each condition described in **H**/**I**. n=3. All values in the graphs are presented as mean±s.e.m. *p≤0.05, ***p≤0.001, ****p ≤0.0001, and ns, non-significant.

Next, we evaluated the status of ER stress in DUSP28 knockdown cells, as it plays a role in the localization of DUSP28 to the mitochondria. The ER morphology in the HeLa cells is a combination of sheets and tubules and has been reported to change their ratios under various physiological conditions (Borgese et al., 2006; Schwarz and Blower, 2016; Shibata et al., 2006). We analyzed the ER morphology in the control and DUSP28 depleted cells by expressing KDEL-RFP (**Fig. 4C**). Interestingly, the number of cells displaying expanded tubular ER morphology was significantly increased and showed enhanced fluorescence intensity of KDEL-RFP in DUSP28 knockdown compared to control cells (**Fig. 4D**). These results suggest an expansion of ER upon knockdown of DUSP28 in HeLa cells. An expanded ER morphology has been reported in cells encountering ER stress to accommodate more chaperones and secretory/transmembrane proteins to combat the stress (Schuck et al., 2009). We investigated if the expansion of ER was due to its stress or any other affected cellular process in DUSP28 knockdown cells. We tested the activation of individual UPR arms by examining the nuclear localization of their downstream transcription factors ATF4, ATF6, and XBP1(s) corresponding to PERK, ATF6 and IRE1 activation, respectively. These transcription factors were reported to translocate to the nucleus upon sensing ER stress (Hetz and Glimcher, 2009; Walter and Ron, 2011). Surprisingly, ATF4 alone but not ATF6 and XBP1(s) showed its enrichment to the nucleus in siDUSP28 compared to siControl cells (**Fig. 4E**). The fluorescence intensity of ATF4 in the nucleus versus total cell was drastically increased in DUSP28 knockdown compared to control cells (**Fig. 4F**). Further, transcriptional and immunoblotting analyses showed enhanced levels of ATF4 in DUSP28 depleted compared to control cells (**Figs. S3J** and **S3K**). These results indicate the activation of the PERK arm of UPR in siDUSP28 cells compared to siControl cells.

Recent studies have shown the contacts between ER and mitochondria and their organelle crosstalk in regulating their function, morphology and biogenesis (Abrisch et al., 2020; Csordas et al., 2018). In line with these studies, the PERK arm of UPR has been shown to regulate mitofusins, which causes the elongation of mitochondria (Lebeau et al., 2018). Consistently, treatment of HeLa cells with ER stress inducer tunicamycin increased the AR of mitochondria (**Figs. 4E** and **4G**). Knockdown of DUSP28 in HeLa cells further enhanced the AR of mitochondria compared to control cells (**Figs. 4E** and **4G**). These studies indicate that the ER stress generated in siDUSP28 cells possibly contributes to mitochondrial expansion (**Fig. 4G**). To test the role of the PERK arm during this process, we treated the siDUSP28 cells with PERKi, which did not restore the AR of mitochondria to the basal level (**Figs. S3L)**. Moreover, transcript levels of different mitochondrial proteins and the protein levels of Mff and Mfn2 were unaffected in siDUSP28 compared to siControl cells (**Figs. S3M** and **S3N**). Thus, the elongation of mitochondria observed in siDUSP28 cells may not be through PERK-dependent signaling. Alternatively, these studies also indicate that the elongation may be due to reduced fission rather than increased fusion since the mitofusin levels are unaffected in siDUPS28 compared to siControl cells (**Figs. S3M** and **S3N**).

The first step in mitochondrial fission involves wrapping ER around mitochondria to form a constriction, which acts as a marking site for mitochondrial fission (Friedman et al., 2011). We tested whether DUSP28 depletion causes a defect in the marking of ER on mitochondria. The live and fixed cell imaging analyses in HeLa cells showed no defect in the marking of ER cross overs with mitochondria rather enhanced over the time of live imaging in DUSP28 knockdown compared to control cells (**Fig. S3C, S3O** and **S3P**). These results indicate that the defects in mitochondrial fission observed in siDUSP28 cells may not be at the initial steps of the process (see below). To test the specificity of DUSP28 in regulating mitochondrial dynamics, we performed a rescue experiment in DUSP28-GFP stable cells, which are transfected with 3’-UTR specific DUSP28 siRNA (**Figs. 4H** and **4I**). As expected, the mitochondrial AR was not enhanced in DUSP28-GFP stable cells upon endogenous knockdown of DUSP28 compared to control HeLa cells (**Fig. 4J**). Hence, DUSP28 plays a role in maintaining the mitochondrial dynamics. We also tested the effect of DUSP28 knockdown on other intracellular organelles in HeLa cells. IFM studies showed an increased number of EEA1- and LAMP1-positive compartments; however, the size was reduced only in LAMP-1-positive organelles in siDUSP28 compared to siControl cells (**Fig. S3Q**, quantified in **S3R-S3U**), indicating a possible indirect role of DUSP28 in regulating the dynamics of these organelles.

### DUSP28 facilitates the dephosphorylation of Drp1 at S637 and causes the mitochondrial fission

Drp1 GTPase mediates mitochondrial fission (Otera and Mihara, 2011) upon its recruitment at the site of constriction marked by the ER (Friedman et al., 2011; Ji et al., 2015). Studies have shown that mitochondrial receptors-Fis1, MiD49, MiD51 and Mff facilitate the recruitment of Drp1 (Loson et al., 2013) to the fission site, following oligomerization, GTP hydrolysis and then cause mitochondrial fission (Frohlich et al., 2013; Smirnova et al., 2001; Yu et al., 2021). The phosphorylation status of Drp1 plays a key in deciding the fission event of mitochondria (Kar et al., 2017). Furthermore, Drp1 has been reported to be phosphorylated at several sites (Elgass et al., 2013), but the phosphorylation status at S^616^ and S^637^ is known to be variable and correlate to the status of mitochondrial fission (Ko et al., 2016). Correspondingly, enhanced levels of phospho-Drp1 (p-Drp1) at Ser^616^ indicate the upregulated mitochondrial fission, while p-Drp1 at Ser^637^ represent the inhibition of fission (van der Bliek et al., 2013). To know the status of p-Drp1 levels in DUSP28 knockdown cells, we performed immunoblotting and measured the levels of total Drp1, p-Drp1(S^616^) and p-Drp1(S^637^) and then compared with the control cells. Surprisingly, siDUSP28 cells showed upregulated levels of p-Drp1(S^637^) as well as total Drp1 without altering the levels of pDrp1(S^616^) in comparison to siControl cells (**Fig. 5A** and **5B**). These results align with the elongated mitochondria observed in siDUSP28 cells due to reduced mitochondrial fission. To validate the role of p-Drp1(S^637^) in siDUSP28 cells, we used phospho-mimetic Drp1^S637D^ and phospho-dead Drp1^S637A^ mutants of Drp1 (Cereghetti et al., 2008). Expression of phospho-dead mutant Drp1^S637A^ displayed shorter mitochondria compared to the Drp1^S637D^ mutant expression, which was similar to the depletion of DUSP28 in HeLa cells (**Fig. 5C**). These studies indicate the dependency of both DUSP28 and phosphorylation status of Drp1 in mediating the mitochondrial fission.

**Figure 5.**
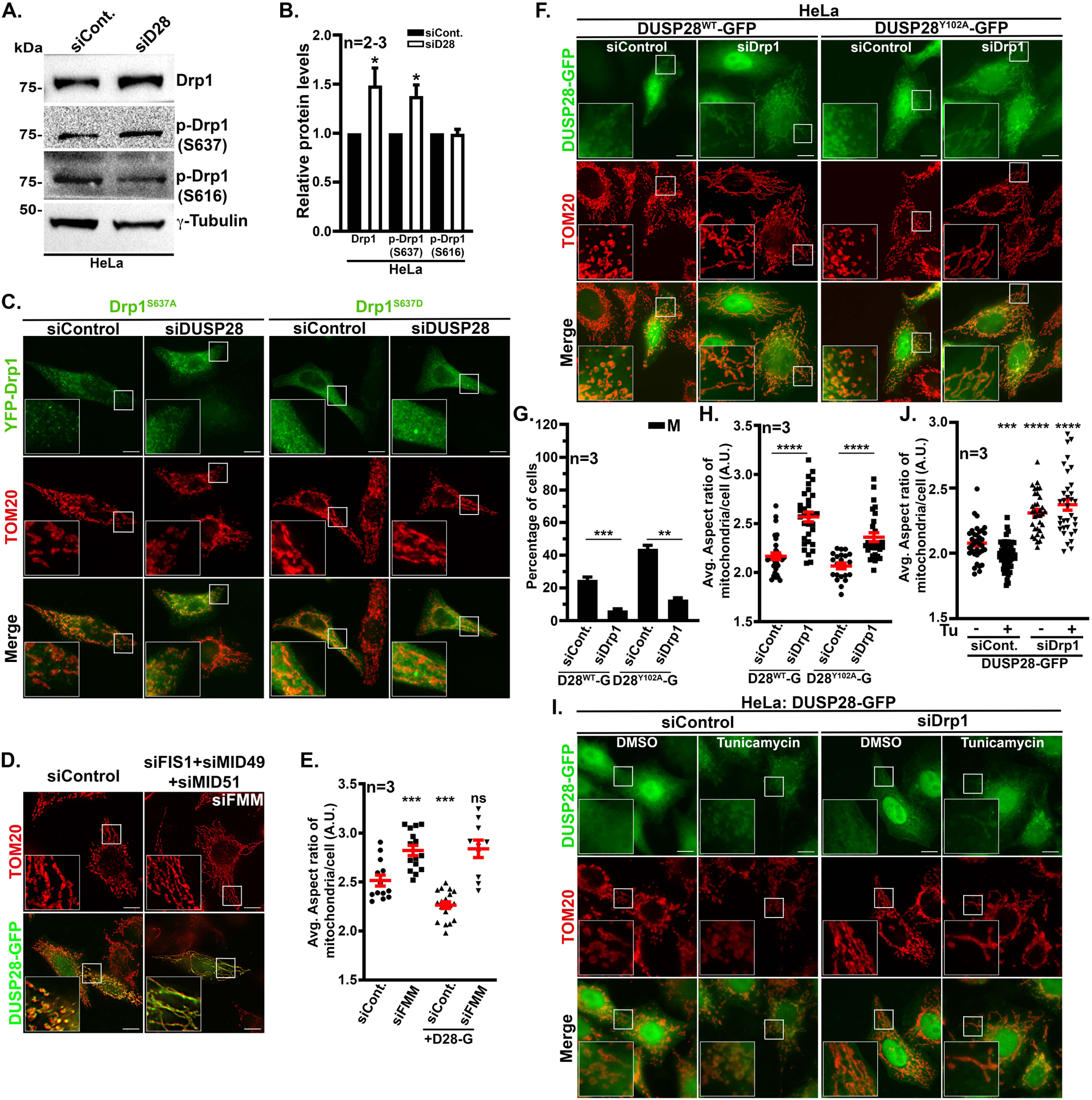
DUSP28 regulates mitochondrial dynamics through Drp1 phosphorylation. **(A)** Immunoblotting analysis of Drp1 protein levels and its phosphorylation status at S637 (p-Drp1(S637)) or S616 (p-Drp1(S616)) in control and DUSP28-knockdown HeLa cells. γ-tubulin is used as a loading control. The relative protein levels for the same were quantified and presented as a graph in **B** (n=3 for Drp1 and p-Drp1(S637), and n=2 for p-Drp1(S616))**. (C)** IFM images of control and DUSP28 knockdown cells that were transfected with YFP-Drp1^S637A^ or YFP-Drp1^S637D^ and stained for TOM20. **(D)** IFM images of control and Fis1+MiD49+MiD51 (siFMM) triple knockdown cells that were transfected with DUSP28-GFP and then stained for TOM20. **(E)** The average aspect ratio (arbitrary units, A.U.) of mitochondria per cell in each condition shown in **D** was quantified and plotted as a graph. n=3. **(F)** IFM images of control and Drp1 depleted cells that were transfected with DUSP28^WT^-GFP or DUSP28^Y102A^-GFP and stained with anti-TOM20 antibody. **(G-H)** Graphs representing the quantification of the percentage of cells with mitochondrial localization of DUSP28-GFP in each condition **(G)** and average aspect ratio (arbitrary units, A.U.) of mitochondria per cell in control and Drp1 knockdown cells with the expression of DUSP28^WT^-GFP (D28^WT^-G) or DUSP28^Y102A^-GFP (D28^Y102A^-G) **(H)**. **(J)** IFM images of control and Drp1 depleted cells that were transfected with DUSP28^WT^-GFP and treated with tunicamycin (Tu) or DMSO. All insets of IFM images are amplified white box areas. Scale 10 μm. **(I)** Graph representing the average aspect ratio (arbitrary units, A.U.) of mitochondria per cell in control and Drp1 knockdown cells treated with DMSO or tunicamycin (Tu). All values are presented as mean±s.e.m. *p≤0.05, **p≤0.01, ***p≤0.001, ****p≤0.0001 and, ns, non-significant.

Studies have shown that the recruitment of Drp1 to mitochondria is facilitated by its recruiters Fis1, MiD49 and MiD51 (Palmer et al., 2011; Yoon et al., 2003). We tested whether these recruiters also regulate the localization of DUSP28 to the mitochondria. To avoid the compensatory function between these molecules, we performed a combined knockdown of Fis1, MiD49 and MiD51 in HeLa cells (referred to here as siFMM cells, **Fig. S4A**) and then expressed DUSP28-GFP. IFM studies showed that the localization of DUSP28-GFP to the mitochondria was unaffected in triple knockdown cells of Fis1, MiD49 and MiD51 (**Fig. 5D**). Further, the mitochondrial localized DUSP28 was unable to cause the fission (**Fig. 5D**), suggesting the requirement of Drp1 and its upstream recruiters Fis1, MiD49 and MiD51 in the DUSP28 mediated mitochondrial fission. Quantification of mitochondrial AR in these experiments (**Fig. 5E**) suggests that DUSP28 recruitment to mitochondria is sufficient to cause mitochondrial fission in siControl cells.

To test the crosstalk or dependency of DUSP28 on Drp1 in regulating mitochondrial fission, we performed siRNA mediated knockdown of Drp1 in HeLa cells and observed the localization of DUSP28^WT^-GFP to mitochondria (**Fig. 5F**). Unexpectedly, depletion of Drp1 significantly reduced the population of cells displaying mitochondrial localized DUSP28 (**Fig. 5G**) and concomitant increase in mitochondrial aspect ratio (**Fig. 5H**). Further, treatment of cells with tunicamycin translocated the DUSP28-GFP to mitochondria but did not cause the fission in siDrp1 cells (**Fig. 5I**). Quantification of mitochondrial AR showed no major difference in morphology upon treatment of DUSP28-GFP expressing siDrp1 cells with tunicamycin (**Fig. 5J**). Additionally, we observed reduced Drp1 fluorescence intensity in cells expressing DUSP28-GFP on mitochondria (**Figs. S4B** and **S4C**), suggesting a crosstalk between these molecules. Next, we used a functional mutant of DUSP28^Y102A^-GFP (described above), which localizes completely to mitochondria and causes fission. Expression of this mutant in Drp1 knockdown cells also inhibited its localization to mitochondria (**Fig. 5G**). Additionally, visualization of a minor cohort of cells expressing DUSP28-GFP (WT and Y102A mutant) on mitochondria did not display mitochondrial fragmentation (**Figs. 5F** and **5H**). These studies emphasize that the fission of mitochondria observed upon translocation of DUSP28 is a Drp1-dependent process. Moreover, the status of p-Drp1(S^637^) may also contribute to the translocation of DUSP28 to mitochondria. Based on these results, we hypothesize that DUSP28 phosphatase modulates the phosphorylation status of Drp1 at S^637^ to regulate the mitochondrial fission.

### DUSP28 inhibits parkin-mediated clearance of damaged mitochondria

As noted, ER stress, but not mitochondrial stress, mediates the translocation of DUSP28-GFP to mitochondria. Here, we tested the role of DUSP28 during the induction of mitochondrial stress. We used CCCP, an oxidative phosphorylation uncoupler that reduces mitochondrial membrane potential, to generate mitochondrial stress (Padman et al., 2013). CCCP is known to affect mitochondrial morphology dramatically and enhances the clearance of damaged mitochondria by targeting to lysosome mediated degradation, called mitophagy (Ashrafi and Schwarz, 2013). mCherry-Parkin has been used as a marker to label the damaged mitochondria, which will be recruited from the cytosol upon sensing mitochondria damage after being phosphorylated by the mitochondrial kinase PINK1 (Kondapalli et al., 2012; Narendra et al., 2010; Vives-Bauza et al., 2010). We tested the localization of DUSP28-GFP and mCherry-Parkin post treatment of HeLa cells with CCCP (10 μM for 6 h). To our surprise, mCherry-Parkin was localized to mitochondria only in cells expressing a cytosolic form of DUSP28-GFP (**Fig. 6A**). In contrast, DUSP28-GFP was observed on mitochondria in a cohort of cells in which mCherry-Parkin localized to the cytosol and was not observed on mitochondria (data not shown). To confirm these results, we used DUSP28^Y102A^-GFP and DUSP28^Y102E^-GFP mutants, which localize majorly to mitochondria and cytosol, respectively. As expected, cells expressing DUSP28^Y102A^-GFP mutant displayed cytosolic localization of mCherry-Parkin, whereas mCherry-Parkin was on mitochondria in cells expressing DUSP28^Y102E^-GFP mutant upon treatment with CCCP (**Fig. 6A**). Based on these results, we hypothesize that the localization of DUSP28 to mitochondria during its damage prevents the recruitment of Parkin possibly by modulating PINK1/Parkin mitophagy. Moreover, we assumed an enhancement of PINK1/Parkin mitophagy in DUSP28 depleted cells during mitochondrial damage. To validate this hypothesis, we induced mitophagy through CCCP in siControl and siDUSP28 cells and studied the mitochondrial clearance. IFM studies showed the localization of mCherry-Parkin to mitochondria (labelled with TOM20), and a cohort of them was targeted to lysosomes (LAMP1-positive organelles) in siControl cells upon treatment of CCCP (**Fig. 6B**). In contrast, the intensity of TOM20 was drastically diminished in CCCP treated siDUSP28 cells and the mitochondria were positive for mCherry-Parkin and LAMP1 (**Fig. 6B**). Quantification of colocalization by measuring the Pearson’s correlation coefficient between the markers confirm the enhanced targeting of mitochondria to lysosomes in siDUSP28 cells during mitochondrial damage (**Fig. 6C** and **6D**). Additional confirmation was obtained using the mito-RFP-GFP construct (Kim et al., 2013), where red puncta would indicate active mitophagy. IFM analysis showed an increase in the number of red puncta in siDUSP28 cells compared to siControl cells (**Figs. 6E** and **6F**). It is important to note that the fission mediated during mitochondrial stress (CCCP-induced) is independent of DUSP28 (**Fig. 6B**). These results indicate that DUSP28 expression on mitochondria inhibits the recruitment of Parkin, thereby inhibiting Parkin-mediated mitophagy.

**Figure 6.**
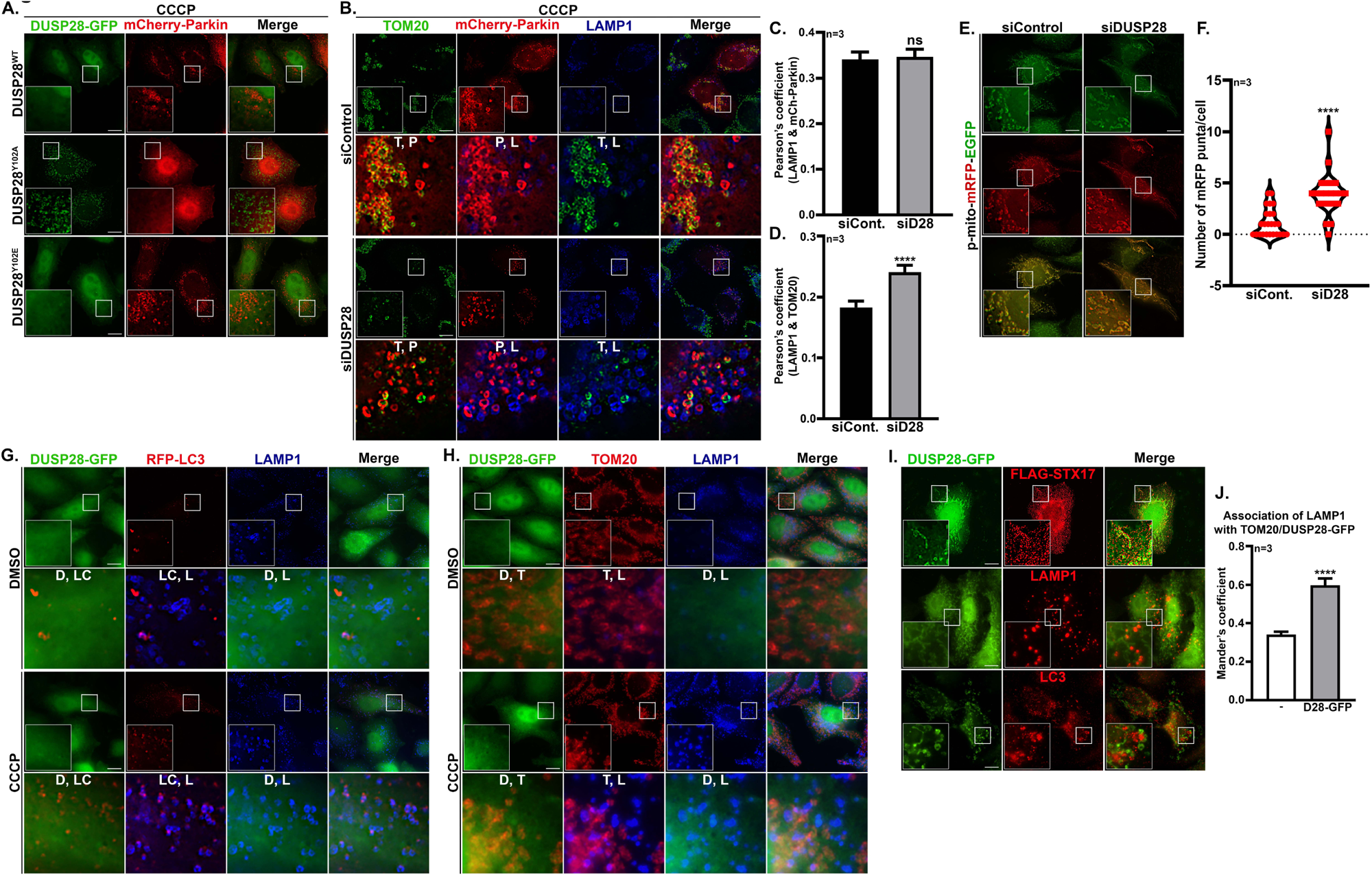
Mitochondrial DUSP28 inhibits Parkin-mediated mitophagy during CCCP induced mitochondrial damage. **(A)** IFM images of HeLa cells co-transfected with DUSP28^WT^-GFP or its mutants DUSP28^Y102A^-GFP or DUSP28^Y102E^-GFP and mCherry-Parkin. Cells were treated with CCCP (10 μm for 6 h). **(B)** IFM images of control and DUSP28 depleted cells that were transfected with mCherry-Parkin. Cells were treated with CCCP (10 μm for 6 h), fixed, and stained with anti-TOM20 and anti-LAMP1 antibodies. **(C-D)** Graphs representing the Pearson’s coefficient values for the colocalization of LAMP1 (L) and mCherry-Parkin (P) **(C)** and LAMP1 (L) and TOM20 (T) **(D)** separately. n=3. **(E)** IFM images of control and DUSP28 depleted cells that were transfected with p-mito-mRFP-EGFP. **(F)** Graph representing the number of mRFP puncta per cell of the similar images shown in **E**. n=3. **(G)** IFM images of stable HeLa cells expressing DUSP28-GFP (D) that were transfected with RFP-LC3 (LC). Cells were treated with DMSO or CCCP, fixed and stained for LAMP1 (L, blue). **(H)** IFM images of stable HeLa cells expressing DUSP28-GFP (D) that were treated with DMSO or CCCP, fixed and stained for TOM20 (T) and LAMP1 (L, blue). **(I)** IFM images of HeLa cells that were transfected with FLAG-STX17 or stained for LAMP1 or LC3 separately. All insets of IFM images are amplified white box areas. In **B, G, H,** insets are enlarged to represent the combination of different colocalization between the proteins. Scale 10 μm. **(J)** Graph represents Mander’s coefficient to show a comparison of LAMP1 association with mitochondria (TOM20) in untransfected and DUSP28-GFP (D28-GFP) in transfected cells. All values are presented as mean±s.e.m. n=3. ****p≤0.0001 and ns, non-significant.

Although DUSP28-GFP in HeLa stable cells did not translocate to mitochondria upon induction of mitochondrial stress, we observed some distinct DUSP28-GFP positive punctate structures in the cells (**Fig. 6G**). To characterize these puncta, we transfected the cells with RFP-LC3 and co-stained with an anti-LAMP1 antibody to represent the autophagosomes and lysosomes, respectively, with the view that these might be targeted for degradation. IFM and line-scan plotting of intensity across the spots showed their positivity with LC3 and LAMP1 (**Figs. 6G** and **S5A**). In addition, these DUSP28-GFP puncta were partly positive for mitochondria (TOM20) (**Fig. 6H** and **S5B**), suggesting that DUSP28-GFP positive mitochondria are directed for lysosomal degradation in a Parkin-independent manner. To investigate the alternative pathway to the Parkin-independent mitophagy in DUSP28-GFP expressing cells, we explored the possibility of STX17-mediated mitophagy (Sugo et al., 2018). Recent studies have shown that STX17 localizes to ER and accumulates onto mitochondria in the Fis1 depletion condition, which directly facilitates the fusion of LC3-positive mitochondria with lysosomes (Xian et al., 2019). We conducted the individual knockdown of Fis1 and Mid49 in HeLa cells and observed the translocation of DUSP28 to mitochondria in siFIS1 but not in siMID49 cells (**Fig. S5C**). Because the localization of DUSP28 to mitochondria requires PERK-mediated ER stress, the status of PERK activation was tested in siFIS1 cells. IFM studies showed the translocation of PERK dependent transcription factor ATF4 to the nucleus only in siFIS1 but not in siControl or siMID49 cells (**Fig. S5D**). Thus, the knockdown of Fis1 in HeLa cells targets DUSP28 to mitochondria via PERK activation, which should also accumulate STX17 on mitochondria. To test this possibility, HeLa cells were transfected with DUSP28-GFP and FLAG-STX17, and their localization pattern was observed. STX17 localization by IFM studies showed the pattern resembling ER tubular morphology in cells expressing FLAG-STX17 alone or with cytosolic DUSP28-GFP (**Fig. S5E**). STX17 appeared as distinct tubular and sheet-like structures in cells expressing mitochondrially localized DUSP28-GFP (**Fig. 6I**). Moreover, FLAG-STX17 showed a significant association with DUSP28-GFP on mitochondria (**Fig. 6I**). As DUSP28-GFP expression on mitochondria drives the accumulation of STX17 to mitochondria, we predict that lysosomes or autophagosomes may follow the same pattern. We observed LAMP1- and LC3-positive structures are also aggregated in the vicinity of DUP28-GFP positive mitochondria (**Fig. 6I**). Consistently, LAMP1 showed enhanced association with mitochondrial localized DUSP28-GFP compared to mitochondrial TOM20 in control HeLa cells (**Fig. 6J**). In line with these results, DUSP28 depletion in HeLa cells mildly altered the levels of STX17 compared to control cells (**Fig. S5F**). Based on these observations, we hypothesize that DUSP28-GFP might initiate the STX17-mediated mitophagy by inhibiting the parkin-dependent mitophagy during mitochondrial damage, which requires future investigation.

## Discussion

DUSP28 belongs to an atypical class of dual specific phosphatase, which dephosphorylates both serine/threonine and tyrosine residues of proteins and non-protein substrates (Pulido and Lang, 2019). Our study showed that DUSP28 localizes majorly to the cytosol and translocates to mitochondria via PERK-dependent ER stress activation. This process regulates the mitochondrial dynamics, an important cellular stress combat pathway to maintain homeostasis. Previous studies connecting the ER stress and mitochondria have shown a significant increase in the mitochondrial length in the presence of ER stress inducers (Lebeau et al., 2018), but our study demonstrated that expression levels of DUSP28 favour the enhanced mitochondrial fragmentation through Drp1-dependent fission. Further, DUSP28 on mitochondria inhibits Parkin recruitment and mediates STX17-dependent mitophagy at the basal level. Thus, these studies provide a key molecular link between ER stress and mitochondrial dynamics controlled by DUSP28 that crosstalks with PERK signaling and Drp1 in maintaining mitochondrial homeostasis, which is critical for cell survival.

DUSP28 localizes as few punctate structures in addition to cytosol at normal conditions. These DUSP28 structures were enhanced during CCCP-induced mitochondrial damage and positive for mitochondria (TOM20), LC3 and lysosomes, indicating the enhanced mitophagy process through STX17 (**Figs. 6 and S5**). However, we predict that this process is highly dependent on expression levels of DUSP28 and PERK activation. Additionally, we observed that DUSP28’s presence on mitochondria (occurs only through ER stress) prevents Parkin-dependent mitophagy during mitochondrial damage. Based on these results, we hypothesize that damaged mitochondria at the basal level may follow DUSP28-STX17 mediated mitophagy rather than Parkin-dependent mitophagy, and the mechanism behind this process requires future investigation. Additionally, this process may support the mitochondrial homeostasis during basal or induced ER stress; however, chemically induced ER stress (tunicamycin, thapsigargin, DTT) requires higher expression of DUSP28. We predict that this might be one of the reasons that the transcript levels of DUSP28 are upregulated in a variety of cancers (Buiga et al., 2018; Lee et al., 2015; Wang et al., 2014) to maintain mitochondrial homeostasis. Interestingly, we observed that cells try to maintain the basal levels of DUSP28 since we lost the expression of DUSP28 in stable cells expressing DUSP28-GFP over a period of time. Further, DUSP28 knockdown cells did not survive during their stable cell preparation using shRNA or CRISPR constructs (data not shown). Thus, DUSP28 play a key role in maintaining mitochondrial dynamics during ER stress.

At the mechanistic level, the function of DUSP28 appears to be dependent on its phosphorylation status both at the active site residue Y102 and outside of the active site residue S125, and their phosphorylation makes the protein cytosol (**Fig. 7**). Our data suggests that these phosphorylation marks will be removed during PERK activation and allow the DUSP28 translocation to mitochondria (see below). It is interesting to study how the conformation changes in DUSP28 allow its interaction with the outer membrane of mitochondria in the future. Additionally, we observed that active site residues (size and phosphorylation status of Y102) play an important role in targeting DUSP28 to mitochondria following its fission (**Fig. 3**). Inline, our mutagenesis analysis of active site residues in DUSP28 justifies its uniqueness in having tyrosine instead of histidine present in other DUSP family members (Ku et al., 2017). For example, replacing tyrosine with histidine (Y102H) in the active site makes the DUSP28 insensitive to ER stress, similar to the reduced phosphatase activity of this mutant in an in vitro assay using purified protein. Further, mutating catalytic residue cysteine to serine in the active site resulted in reduced/no activity, alike the previous *in-vitro* results (Ku et al., 2017). Thus, tyrosine in the active site does play a critical role in regulating DUSP28 translocation to mitochondria in mediating fission; possibly being phosphorylated will cause the protein to be in the cytosol. While studying PERK signaling and its downstream phosphatase calcineurin linking to DUSP28, we identified another residue, S125, that exhibited a major role in deciding the status of DUSP28 localization. Our data suggests that phospho-S125 in DUSP28 is possibly dephosphorylated by calcineurin during PERK activation, which may cause a conformation change to expose active site phospho-Y102 for further dephosphorylation before translocating to mitochondria (**Fig. 7**). Further, the association of DUSP28 to outer membrane of mitochondria possibly occurs through an unknown mitochondrial receptor as the depletion of known receptors (Fis1, MiD49, MiD51) showed no effect on DUSP28 localization (**Figs. 5 and S4**).

**Figure 7.**
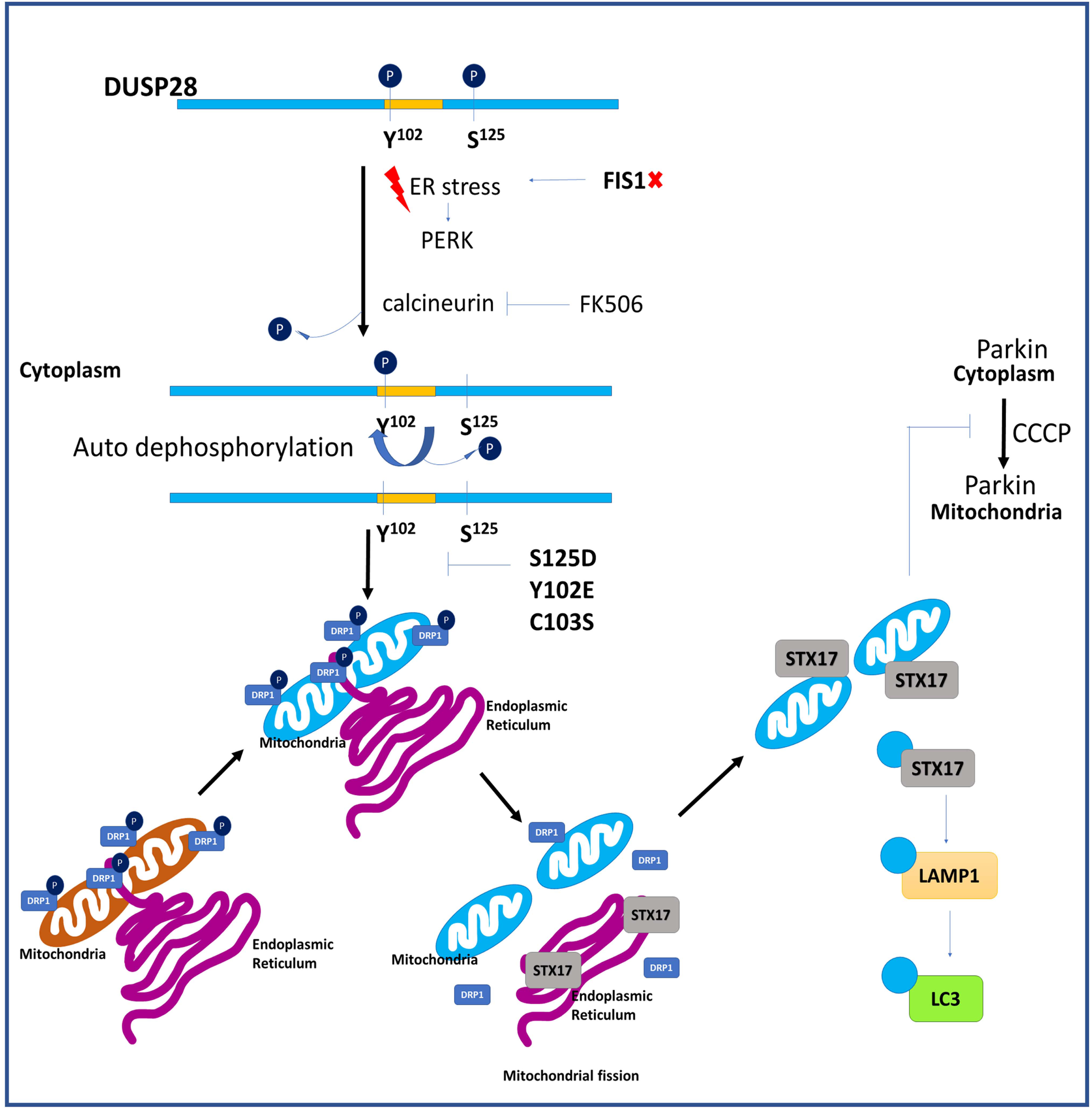
Schematic model representing the mechanism of DUSP28 localization to mitochondria and its regulation on mitochondrial dynamics and turnover. DUSP28 sequence analysis using NetPhos 3.1 (DTU Health Tech, Denmark) identified two predicted phosphorylation sites at Y102 (in the active site) and S125, which are responsible for its cytosolic localization. During ER stress, the phosphatase calcineurin possibly dephosphorylates DUSP28 at the S125 position, which can be inhibited by GSK2606414 (PERK inhibitor) or FK506 (calcineurin inhibitor). Further, an unknown phosphatase or auto dephosphorylation at the Y102 position causes DUSP28 translocation to mitochondria. This step also dependents on the phosphorylation status of S125, as DUSP28 (S125D/Y102E/C103S) mutants block its translocation to mitochondria. Upon translocation to mitochondria, DUSP28 dephosphorylates Drp1 at S637 and promotes mitochondrial fission. The recruitment of DUSP28 to mitochondria prevents Parkin localization during the damage induced by CCCP and follows STX17-mediated Parkin-independent mitophagy. The STX17-DUSP28 positive mitochondria recruit LC3 and are targeted for lysosomal degradation.

DUSP28 recruitment to the mitochondria upon PERK activation possibly interacts and dephosphorylates S637 of Drp1, which promotes mitochondrial fission. Consistently, Drp1^S637A^ but not Drp1^S637D^ mutant rescued the mitochondrial fragmentation in DUSP28 depleted cells (**Fig. 5**). To our surprise, depletion of Drp1 reduces the mitochondrial translocation of DUSP28, suggesting a crosstalk between these molecules during fission. We further hypothesize that the DUSP28 possibly interacts with p-Drp1 brought by the ER at the mitochondrial fission sites. Post-dephosphorylation of Drp1 by DUSP28 may facilitate mitochondrial fission during ER stress. However, mitochondrial damage induced by CCCP does not affect the recruitment of DUSP28 to mitochondria. The presence of DUSP28 on mitochondria prevented the recruitment of Parkin to mitochondria by an unknown mechanism. This process favoured the STX17-mediated mitophagy to clear the damaged mitochondria upon CCCP treatment of HeLa cells (**Fig. 6**). These studies suggest that DUSP28 critically regulates mitochondrial homeostasis during both ER and mitochondrial stress conditions. Conversely, cells favour faster clearance of mitochondria by Parkin-mediated mitophagy (induced by CCCP) upon DUSP28 knockdown, indicating DUSP28 acts as a switch between Parkin- and STX17-mediated mitophagy during mitochondrial damage.

Based on our results, we proposed a model for the function of dual specificity phosphatase DUSP28 in mitochondrial fission (**Fig. 7**). DUSP28 is primarily localized to the cytosol due to its assumed dual phosphorylation at Y102 and S125 residues. During ER stress following PERK activation, the activated calcineurin dephosphorylates S125 of DUSP28, possibly allowing the exposure of phosphorylated Y102 residue. We hypothesize that DUSP28 removes the phosphorylation mark at Y102 through self-auto-dephosphorylation or post dimerization (reported in (Ku et al., 2017), which allows the DUSP28 translocation to mitochondria. Consistently, DUSP28 translocation can be inhibited using inhibitors of the PERK arm GSK2606414 or calcineurin inhibitor FK506. In addition, the active site residue C103 also plays a key role in DUSP28 translocation to mitochondria. DUSP28 on mitochondria facilitates its fragmentation through the conventional Drp1-mediated fission post dephosphorylation of Drp1 at the S637 position. However, the loss of the Drp1 function or its recruiters (such as Fis1, MiD49, MiD51) cannot be rescued with mere overexpression of DUSP28. Thus, we predict that DUSP28 on mitochondria dephosphorylates p-Drp1(S637) brought to the fission site by ER that causes the mitochondrial fission. Interestingly, DUSP28 facilitated the recruitment of STX17 by inhibiting Parkin to the damaged mitochondria (induced by CCCP) and targeting it to mitophagy. This process may occur if DUSP28 dephosphorylates Parkin on mitochondria during its damage, which requires future investigation. In a nutshell, we uncover a novel function for an atypical phosphatase DUSP28 in maintaining mitochondrial dynamics during ER stress.

### Future Perspectives

Our study uncovers a novel role for DUSP28 in connecting various cellular homeostasis pathways, such as mitochondrial health during ER stress. In our time chase experiments, we also observed a cyclic phase of DUSP28 localization to mitochondria, which may correlate to the PERK activation (Malhotra and Kaufman, 2007; Mori, 2009). During the initial phase of ER stress, DUSP28 localizes to mitochondria and maintains homeostasis by blocking Parkin-mediated mitophagy; as stress is prolonged, DUSP28 translocates to cytosol that restores Parkin-dependent mitophagy. This process is possibly adopted by several cancers, wherein DUSP28 transcripts are highly abundant, which maintains the mitochondrial homeostasis required for tumor growth. In the future, structural studies in the context of cellular signaling may help in developing small molecules against DUSP28 and form a therapeutic target. Thus, this study is the first of a kind to connect ER stress and mitochondrial dynamics through DUSP28.

## Methods

### Reagents and antibodies

All general chemicals, including bafilomycin A1, carbonyl cyanide m-chlorophenyl hydrazone (CCCP), ceapin-A7 (ATF6 inhibitor, ATF6i), DMSO, 1,4-dithiothreitol (DTT), FK506 (calcineurin inhibitor), GSK2606414 (PERK inhibitor, PERKi), puromycin, MG132, thapsigargin and tunicamycin were purchased from Sigma-Aldrich (Merck). Rotenone and KIRA6 (IRE1α inhibitor, IRE1i) were purchased from Calbiochem. 10-N-Nonyl acridine orange (NAO), tetramethylrhodamine ethyl ester perchlorate (TMRE), MitoTracker™ Red CMXRos, MitoTacker Green and tissue culture reagents were purchased from Thermo Fisher Scientific (Invitrogen). The following commercial polyclonal and monoclonal antisera were used in this study: anti-ARF1+ARF3 (Ab32524) and anti-DRP1 (Ab56788) were from Abcam; anti-GM130 (610822) and anti-p230 trans Golgi (611281) were from BD Biosciences; anti-ATF4 (11815), anti-BiP (3177), anti-Calnexin (2679), anti-Phospho-DRP1(S616) (4494), anti-Phospho-DRP1(S637) (6319), anti-EEA1 (3288), anti-IRE1 (3294), anti-MFF (84580), anti-Mitofusin-2 (11925), anti-Parkin (4211) and anti-PERK (5683) were from Cell Signaling Technology; anti-hLAMP-1 (H4A3) was from Developmental Studies Hybridoma Bank; anti-FIS1 (10956-1-AP) was from ProteinTech; anti-ATF-6α (sc-22799), anti-GAPDH (sc-25778), anti-Tom20 (sc-17764) and anti-XBP-1 (sc-7160) were from Santa Cruz Biotechnology; anti-β-Actin (A5441), anti-LC3B (L7543), and anti-γ-Tubulin (T6557, IB:1:10000) from Sigma-Aldrich (Merck); and anti-GFP (A-11122) was from Thermo Fisher Scientific (Invitrogen). All secondary antibodies were either from Invitrogen or Jackson ImmunoResearch. All primary antibodies were used at 1:100 dilution for IFM and 1:1000 dilution for immunoblotting. The secondary antibodies were used at 1:500 dilution for IFM and 1:10000 dilution for immunoblotting.

### siRNA sequences

The following control siRNA (siControl) and target siRNA sequences against the respective gene were synthesized from Eurogentec, Belgium. siDUSP28 (5’-CUGGUCUCAGCUCCAGAAGUAUG-3’), siDUSP28-3’-UTR (5’-CGUGUCCCAUGAAUCUUGU-3’), siDrp1 (5’-UGCUAUGGUAAUGCACUUG-3’), siFIS1 (5’-AACGAGCUGGUGUCUGUGGAG-3’), siMID49 (5’-ACUUUCGGAGCAAGUUCCCGGAACU-3’) and siMID51 (5’-GCCAAGCAAGCUGCUGUGGACAUAU-3’).

### Primers used for transcript analysis

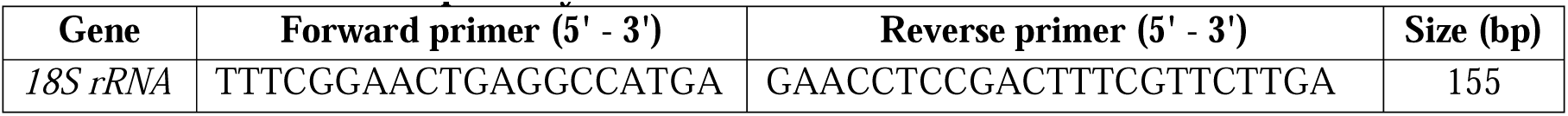
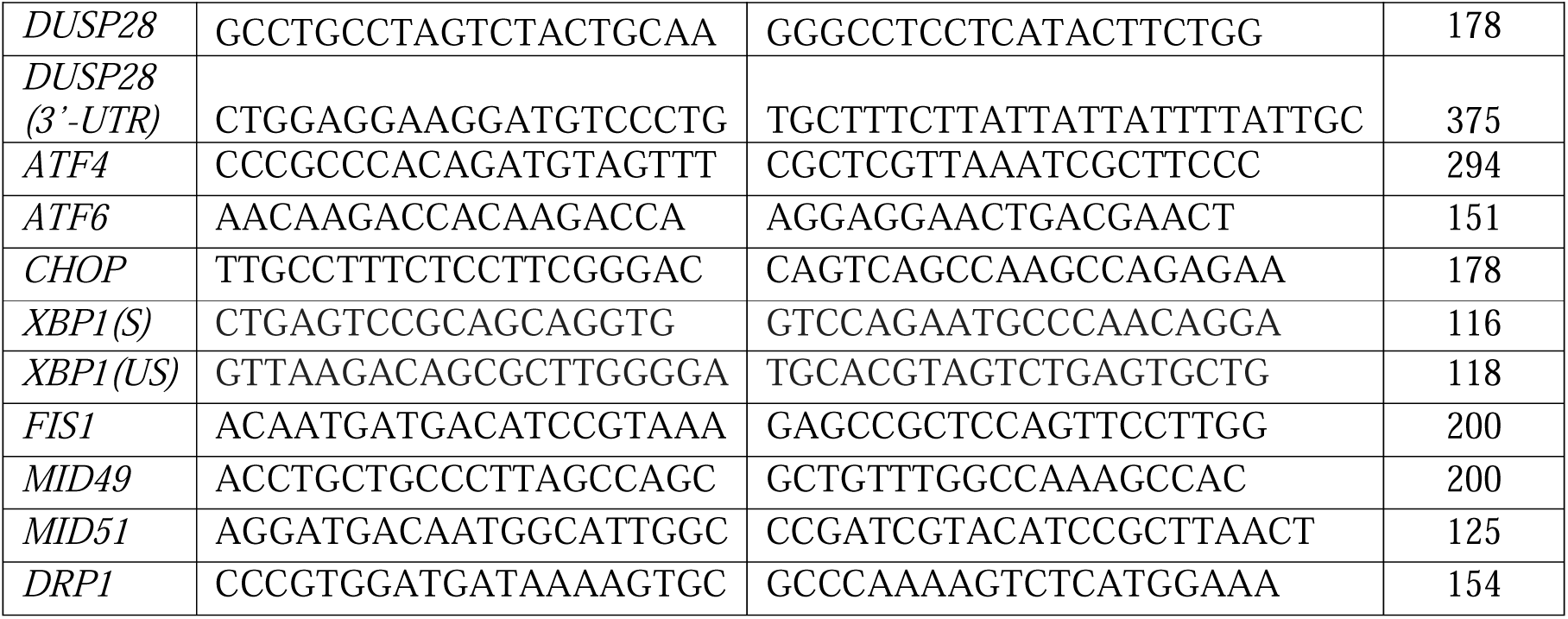

### DNA constructs

#### pEGFP-N3-DUSP28

DUSP28 gene was PCR amplified using gene specific primers from the cDNA, prepared from HeLa cell lysate. The PCR product was digested and cloned at *Bgl*II and *Sal*I sites of the pEGFP-N3 vector. The clones of the DUSP28 plasmid were confirmed by restriction digestion followed by DNA sequencing. Analysis of the DNA sequence showed two additional mutations contributing to a change in amino acids at S52P and C99R compared to the reported sequence NP_001357394.1 in the NCBI database. These mutations are also present in the cDNA that was amplified from HEK293T cell lysate. The following DUSP28-GFP mutants were generated through site-directed mutagenesis (Shakya et al., 2018) using pEGFP-N3-DUSP28 (WT) as a plasmid template: DUSP28(P52S)-GFP, DUSP28(P52S,R99C)-GFP, DUSP28(Y102H)-GFP, DUSP28(Y102A)-GFP, DUSP28(Y102E)-GFP, DUSP28(Y102F)-GFP, DUSP28(C103S)-GFP, DUSP28(S13D)-GFP, DUSP28(S125D)-GFP and DUSP28(S125A)-GFP. All plasmids were sequenced and verified.

#### pmCherry-N3-DUSP28

mCherry gene was PCR amplified from pmCherry-C1 vector and digested with *Bam*H1 and *Not*I enzymes and then replaced with GFP in pEGFP-N3-DUSP28 vector. The clone was verified by restriction digestion followed by DNA sequencing.

#### pEGFP-N3-sDUSP28

sDUSP28 was PCR amplified using pEGFP-N3-DUSP28 as a template, digested with *Bgl*II and *Sal*I restriction enzymes and then cloned in pEGFP-N3 vector at the same sites. The clone was confirmed by restriction digestion and then with DNA sequencing.

#### pLVX-Puro-DUSP28

DUSP28 was PCR amplified using pEGFP-N3-DUSP28 as a template, digested with *Eco*RI and *Xba*I restriction enzymes and then cloned into pLVX-Puro (Clontech) vector at the same sites. The clone was confirmed by restriction digestion followed by DNA sequencing.

##### Other plasmids

YFP-Drp1 (WT), YFP-Drp^S637A^ and YFP-Drp1^S637D^ were obtained as a gift from Prof. Luca Scorrano, Universita degli Studi di Padova, Italy (Cereghetti et al., 2008). mCherry-Drp1 (49152), pMD2.G (12259), psPAX2 (12260), mCherry-Parkin (23956), DsRed2-Mito-7 (55838), pmRFP-LC3 (21075) and FLAG-Stx17 (45911) were obtained from Addgene. KDEL-RFP (Singh et al., 2018) and p-mito-mRFP-EGFP (pAT016) (Kim et al., 2013) were described previously.

### Cell culture, transfection and lentiviral transduction

HeLa and HEK293T were procured from the American Type Culture Collection (ATCC). Cells were maintained in Dulbecco’s modified Eagle’s medium (DMEM, Invitrogen) supplemented with 10% FBS (Biowest), 1% L-glutamine (Invitrogen) and 1% penicillin– streptomycin (Pen-Strep, Invitrogen) antibiotics at 37°C with 10% CO_2_ in a humified cell culture incubator. Transfection of cells with plasmid DNA or siRNA was carried out using Lipofectamine 2000 (Invitrogen) or Oligofectamine (Invitrogen), respectively, according to the manufacturer’s protocol. Lentivirus expressing DUSP28-GFP was prepared using a protocol described previously (Shakya et al., 2018). Briefly, HEK293T cells were seeded in a 35 mm dish pre-coated with Matrigel (BD Biosciences, prepared in 5% serum free DMEM medium). Cells were transfected with pLVX-Puro-DUSP28-GFP plasmid (250 ng) along with pMD2.G (250 ng) and psPAX2 (1000 ng) in a 1:1:4 ratio of DNA. Post 60 h of transfection, the virus was collected, filtered using a 0.45µm filter and then added to the HeLa cells along with polybrene (10 μg/ml). Post 48 h of transduction, cells were selected with a medium containing puromycin gradually (1 to 5 μg/ml) for 15 days. Henceforth, the cells were always maintained in a medium containing 1 µg/ml puromycin. Early passaged stable cells were used for the experiments. In Figs. 2, 3, 4, 5 and 6 and Figs. S2, S5, the cells were treated with different drugs as follows: tunicamycin (Tu, 10 μM for 6 h), FK506 (3 nM for 6 h), thapsigargin (Tg, 5 μM for 6–12 h), rotenone (500 nM for 6 h), bafilomycin A1 (5 mM for 6 h), MG132 (5 μM for 4 h), KIRA6 (1 μM for 12 h), ceapinA-7 (5 μM for 12 h), CCCP (10 μM for 6 h) and GSK2606414 (0.1 μM for 12 h). As a control, DMSO was used during these treatments.

### Immunoblotting

Cells were lysed in RIPA buffer (10 mM Tris-HCl pH 7.4, 5 mM EDTA, 50 mM NaCl and 0.5% v/v NP40) in the presence of protease inhibitor cocktail (SIGMA*FAST*) on ice for 30 min. Homogenates were centrifuged at 10000 rpm for 10 min to remove cell debris, including nuclei. Protein concentration of the lysate was measured using the Bradford reagent according to the manufacturer’s instructions (Bio-Rad Laboratories). Protein extracts were denatured with Laemmli buffer (62.5 mM Tris-Base pH 6.8, 8% glycerol, 2.3% sodium dodecyl sulfate (SDS), 0.005% bromophenol blue and 5% 2-mercaptoethanol) for 5 min at 100°C. Protein amounts equivalent to 50–100 μg were subjected to SDS–polyacrylamide gel electrophoresis (Bio-Rad Laboratories) and then transferred onto polyvinylidene difluoride (PVDF) membranes. Membranes were incubated in a blocking solution (1xPBS, 0.02% Tween-20 and 5% milk) for 1 h at room temperature and then probed with respective primary followed by secondary antibodies. Immunoblots were developed with Clarity Western ECL substrate (Bio-Rad Laboratories) and then imaged in a Bio-Rad Molecular Imager ChemiDoc XRS+ imaging system using Image Lab 4.1 software. Protein band intensities were quantified and then normalized with a loading control. The fold change in values with respect to control is indicated in the figure.

### Immunofluorescence microscopy (IFM)

Cells were seeded in 24 well plate containing glass coverslips at a density of 30% confluence. Cells were treated with compounds as indicated in each experiment. Cells were fixed with 4% paraformaldehyde, blocked with 3% bovine serum albumin (BSA) and stored in 1xPBS. For staining, the probe MitoTracker™ Red CMXRos was added to the cells at 400 nM and incubated for 20 min prior to fixation. Cells on glass coverslips were incubated with primary antibodies for 30 mins at room temperature. Coverslips were washed in 1xPBS and then incubated for 30 mins with anti-mouse or anti-rabbit Alexa-conjugated secondary antibodies. Coverslips were finally washed and mounted on glass slides using Fluoromount-G. IFM imaging of the cells was performed on an Olympus IX81 motorized inverted fluorescence microscope equipped with a CoolSNAP HQ2 (Photometrics) CCD camera using a 60x (oil) U Plan super apochromatic objective. The deconvolution of images (except for the cells with cytosolic signal) was carried out using the cellSens Dimension package with the 5D module (Olympus) and then analyzed. Note that in Fig. 1, the z-stacks were subjected to constrained iterative deconvolution (1 or 2 iterations) using the cellSens Dimension package. Otherwise, undeconvolved z-stack images were converted to maximum intensity projection (MIP) for further analyses/assembly. Images were assembled using Adobe Photoshop.

For live cell imaging, cells were seeded in live cell dishes (MatTek Life Sciences) and then transfected with DUSP28-GFP and mitochondrial marker DsRed2-Mito7 or DUSP28-GFP transfected cells were stained with 100 nM MitoTracker Green in a complete medium for 30 mins at 37°C. Cells were imaged under the Olympus IX81 wide-field microscope, and the movie was captured for live tracking of mitochondria.

### Methods to quantify the images of IFM and cellular assays

In all the methods, processed/MIP images were used for further image analyses.

### I. Measurement of corrected total cell fluorescence (CTCF) intensity

CTCF was used to measure the total cell fluorescence intensity. Briefly, the area of the cell, mean fluorescence intensity and integrated density parameters were measured using ImageJ. Using the freehand selection tool, an individual cell was selected in the field, and then measured the parameters mentioned above (Go to Analyze > measure > give result separate result sheet > copy the results to Excel). Similarly, the data for the background was also calculated for each quantified image. The CTCF was calculated using the formula: CTCF = Integrated density – (Area of the cell * Mean fluorescence of the background). The data was analyzed and plotted as graphs using GraphPad prism 8.4.2.

### II. Measurement of colocalization coefficient values

The Pearson’s correlation coefficient (*r*) value between two fluorophores was estimated using Olympus cellSens Dimension software. The MIP of undeconvolved z-stack images were used for the analysis. Additionally, multiple ROIs (regions of interest) were selected in the cytosol by excluding the perinuclear area. The average *r* value from 20 to 30 cells of each experiment, performed in triplicates or more, was calculated and indicated as mean±s.e.m.

### III. Measurement of Mander’s overlap between the fluorophores

The Mander’s coefficient value between two fluorophores was estimated using JACoP (Just Another Colocalisation Plugin) of ImageJ 1.53 software. The MIP of the desired image was selected, followed by the splitting of channels (Image> Color> Split Channels) and uploaded as Image A and B in the plugin. M1 and M2 coefficient options from the tick box were selected and analyzed. The M1 for a set was analyzed and then plotted using GraphPad prism 8.4.2.

### IV. Quantification of the number of fluorescent puncta

IFM images were opened through the Olympus viewer plugin of ImageJ 1.53 software. The single channel of an image was selected using the split channel tool present under the image tab. The z-stacks were merged into a single plane using the z-projection tool (present within the stacks dropdown) available under the image tab. The selected single channel MIP image was converted to an 8-bit image followed by rolling ball background subtraction (process tab > subtract background) with a 20.0 pixels radius. The image was adjusted to auto thresholding (30 to 255; default; Image tab > adjust > threshold). Following, the image was converted into binary (Process tab > binary > make binary), which succeeded with watershed segmentation (Process tab > binary > watershed). Using the freehand selection tool, within a relative cell area, the number of puncta was calculated for each cell (analyze tab >analyze particles). The puncta count was calculated based on particle size at a default range between 0.0-infinity (μ2) with a circularity between 0.00-1.00. The number of puncta per cell was analyzed and then plotted using GraphPad prism 8.4.2.

### V. Percentage of cells with mitochondrial localization of DUSP28-GFP

HeLa cells were transfected with DUSP28-GFP or its mutants and stained for mitochondria using an anti-TOM20 antibody. Cells were imaged and analyzed in ImageJ 1.53 software. The number of cells showing the DUSP28-GFP or its mutant’s localization to the cytosol, mitochondria or both were counted visually. All quantifications were carried out in a blinded fashion. The percentage of cells with mitochondrial or cytosolic localization was calculated using the formula:

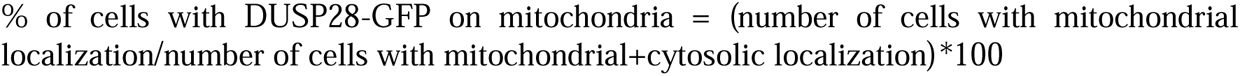

The calculations were performed in Microsoft Excel 10 and then plotted using GraphPad prism 8.4.2.

### VI. Quantification of mitochondria by aspect ratio

Mitochondrial quantification includes 2D analysis of morphology and its network characteristics. For 2D analysis, the image was first analyzed for “Tubeness” (sigma 0.01075) and then converted to an 8-bit image that was thresholded using the “default” option (“Adjust” in the dropdown menu of the “image” option). The resulting binary image was used as input to analyze the mitochondria using the “Mitochondria Analyzer” plugin, and the 2D option was further selected (Chaudhry et al., 2020). The obtained results consist of multiple parameters, including number, aspect ratio, form factor, etc. The average aspect ratio of mitochondria per cell was analyzed and then plotted using GraphPad prism 8.4.2.

### VII. Quantification of the percentage of tubular ER

The Z-stack of a cell was converted into a composite image using Fiji (“Image”→“Stacks”→“Z project”→select slice→OK). Select the following settings beforehand for measuring the area: “Analyze”→“Set Measurements”→“Check the Area”→OK. Then, select the whole cell area by using a freehand tool (Choose freehand selection→select the area→“Analyze”→“Measure”). Further, copy the area number and transfer the data to an Excel file. Similarly, select the area covering the ER sheet and measure the percentage of sheet area (=sheet area/total area * 100). Calculate the percentage of tubular ER using the formula: percentage of tubular area = 100-percentage of sheet area.

### Isolation of mitochondria

Stably expressing DUSP28-GFP HeLa cells were cultured in approximately 6 – 8 number of 10 cm dishes. At around 80% of cell density, the dishes were divided into two sets. One set was treated with DMSO and another set with tunicamycin (10 μM for 6 hrs). After the treatment, the cells were washed twice with 1xPBS and then trypsinized. The obtained pellet was subjected to crude mitochondrial isolation protocol (Wieckowski et al., 2009), which involves the homogenization of the cell pellet in the homogenization buffer (225 mM mannitol, 75 mM sucrose, 0.1 mM EGTA and 30 mM Tris–HCl pH 7.4). The homogenized lysate was centrifuged at 600*g* to remove nuclei and unbroken cells. This step was repeated twice, and the obtained supernatant was subjected to a 10 min spin at 7000*g* to obtain the crude mitochondria. The pellet was washed twice or thrice by suspending the mitochondria in suspension buffer (225 mM mannitol, 75 mM sucrose and 30 mM Tris–HCl pH 7.4) and collected after a final spin of 10,000*g.* The final mitochondrial pellet was resuspended in mitochondria resuspending buffer (MRB, 250 mM mannitol, 5 mM HEPES (pH 7.4) and 0.5 mM EGTA) and then subjected to immunoblotting.

### Statistical analysis

GraphPad Prism 8.4.2 software was used to calculate the statistical significance. Statistical significance was determined by the unpaired Student’s t-test and variance analysis. Ns, not significant; *, p ≤0.05; **, p ≤0.01 and ***, p ≤0.001.

## Acknowledgements

We thank Samit Ravindra Desai for conducting site-directed mutagenesis of DUSP28-GFP constructs. This work was supported by the Department of Biotechnology (BT/PR32489/BRB/10/1786/2019), Science and Engineering Research Board (CRG/2019/000281), DBT-NBACD (BT/HRD-NBA-NWB/38/2019-20), India Alliance (500122/Z/09/Z), and IISc-DBT partnership programme. Infrastructure in the department was supported by DST-FIST, DBT, and UGC. PB was supported by DBT-JRF (DBT/2016/IISc/717).

## Author contributions

PB performed all the experiments described in this study. SRGS designed, oversaw the entire project, coordinated and discussed the work with co-author and wrote the manuscript.

## Supplementary Figures

**Figure S1.**
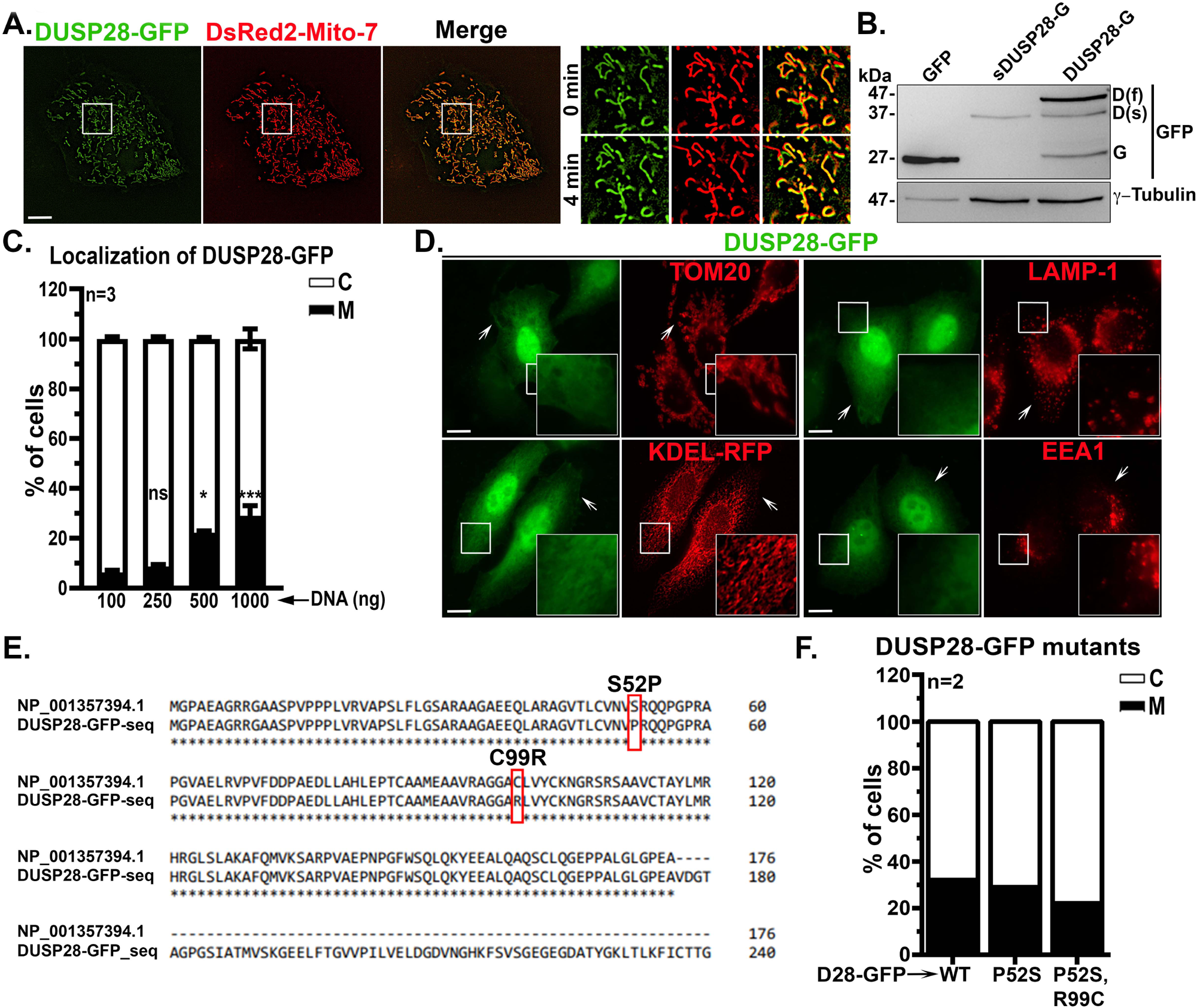
**(A)** Live cell imaging of mitochondrial DUSP28-GFP along with DsRed2-Mito-7. Single frames of a live cell movie in HeLa cells were shown separately. Merged image showing the complete colocalization of membranous DUSP28 with a mitochondrial marker. Insets represent the white boxed area of the cell taken at 0 min and 4 min time points to validate the mitochondrial localization of DUSP28-GFP over the time. **(B)** Immunoblotting analysis of HeLa cell lysates expressing GFP, a shorter isoform of DUSP28 (sDUSP28-GFP) or DUSP28-GFP (DUSP28-G). Blot was probed with anti-GFP antibody. The observed bands correspond to GFP (G; 27 kDa), sDUSP28-GFP(D(s); 37 kDa) and full-length DUSP28-GFP (D(f); 48 kDa). γ-Tubulin is used as a loading control. **(C)** Graph represents the percentage of cells showing the cytosolic (C) or mitochondrial (M) localization of DUSP28-GFP with increased plasmid concentration during the transfection of HeLa cells. All values are presented as mean±s.e.m. n=3. *p≤0.05, ***p≤0.001, and ns=non-significant. **(D)** IFM images of HeLa cells transfected with DUSP28-GFP (cytosolic) and co-stained with the markers of mitochondria (TOM20), lysosomes (LAMP1) and endosomes (EEA1). Cells were transfected with KDEL-RFP to represent ER. Arrows point to the cytosolic DUSP28-GFP. Insets are amplified white box areas. Scale 10 μm. **(E)** Annotated protein sequence of DUSP28 that was cloned from the cDNA of HeLa cells. Sequence was aligned with the reported protein sequence of DUSP28 (NP_001357394.1). Alignment of both the sequences using ClustalW showed two mutations, S52P and C99R, in the cloned DUSP28 sequence. **(C)** Graph represents the percentage of cells showing cytosolic (C) or mitochondrial (M) localization of DUSP28-GFP (D28-GFP) in HeLa cells expressing wildtype (WT), P52S or P52S,R99C mutants of DUSP28. Note that both the mutants of DUSP28-GFP showed equal tunicamycin sensitivity, similar to WT. n=2.

**Figure S2.**
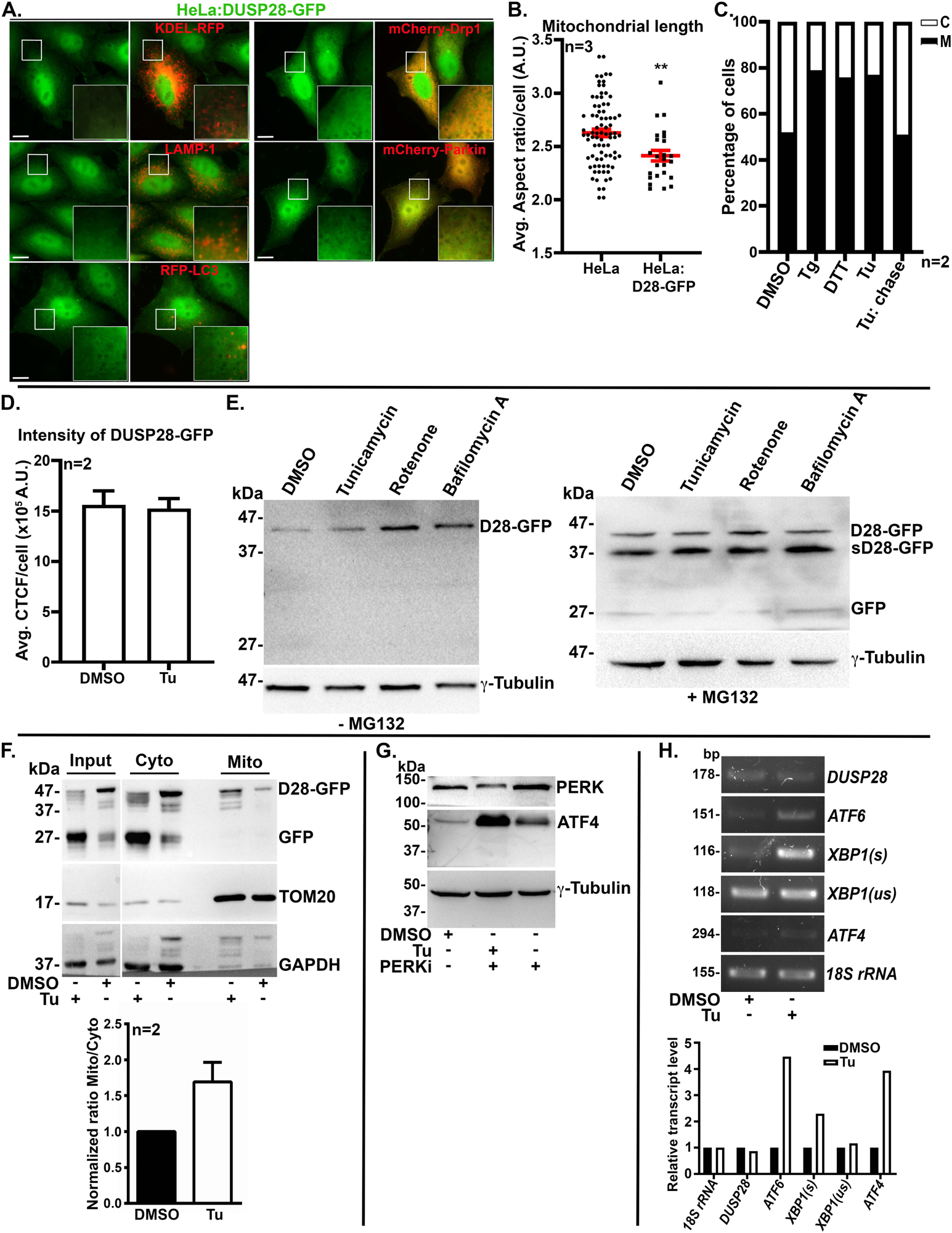
**(A)** IFM images of HeLa cells stably expressing a cytosolic form of DUSP28-GFP (HeLa:D28-GFP), co-stained with anti-LAMP-1 antibody (lysosomes) or transfected with KDEL-RFP (endoplasmic reticulum) or RFP-LC3 (autophagosomes). Cells were transfected with mCherry-Drp1 or mCherry-Parkin to study their localization in the presence of stably expressed DUSP28-GFP. Insets are amplified white box areas. Scale bar, 10 μm. **(B)** Graph represents the aspect ratio (arbitrary units, A.U.) of mitochondria per cell in HeLa and HeLa:DUSP28-GFP stable cells. n=3. **p≤0.01. **(C)** Graph indicates the percentage of cells with mitochondrial or cytosolic localization of DUSP28-GFP after the treatment with various ER stress inducers – thapsigargin (Tg; 500 nM for 6 h), DTT (1 mM for 6 h) or tunicamycin (Tu, 10 μm for 6 h). DMSO is used as a control. The last bar represents the effect on DUSP28-GFP localization upon removal of Tu post 6 h of the chase in a plain medium. n=2. **(D)** Graph represents the fluorescence intensity of DUSP28-GFP (measured by CTCF) in HeLa:D28-GFP stable cells treated with tunicamycin (10 μm for 6 h) or DMSO. N=2. € Immunoblotting analysis of HeLa:D28-GFP lysates. Cells were treated with various organelle’s stress inducers such as tunicamycin, rotenone or bafilomycin A and DMSO as a control in the absence and presence of the proteasomal inhibitor MG132 (5 μM for 4 h). Note that the expression of sDUSP28-GFP was observed in the cell lysates treated with MG132 (right blot). γ-tubulin is used as a loading control. **(F)** Immunoblotting analysis of HeLa:D28-GFP lysates subjected to sub-cellular fractionation to obtain the crude mitochondria and cytosol fractions. Cells were treated with DMSO or tunicamycin before the fractionation (described in Materials and Methods). Graph indicates the quantification of DUSP28-GFP participation to mitochondria vs cytosol fractions (represented as a normalized ratio of D28-GFP intensities in Mito/Cyto). GAPDH is used as the loading control. n=2. All values are presented in the graphs as mean±s.e.m. **(G)** Immunoblotting analysis of lysates of HeLa cells treated with DMSO, tunicamycin or tunicamycin+PERKi (PERKi 100 nM 6 h). Lysates were probed for the expression of PERK and ATF4 to validate the compound treatments. γ-tubulin is used as a loading control. **(H)** Semiquantitative PCR analysis of DUSP28 and UPR genes expression in cell lysates treated with DMSO or tunicamycin (10 μM for 6 h). *18S rRNA* is used as an internal control. s – spliced and us – unspliced forms of XBP1. Graph represents the relative transcript levels calculated based on the gel intensities.

**Figure S3.**
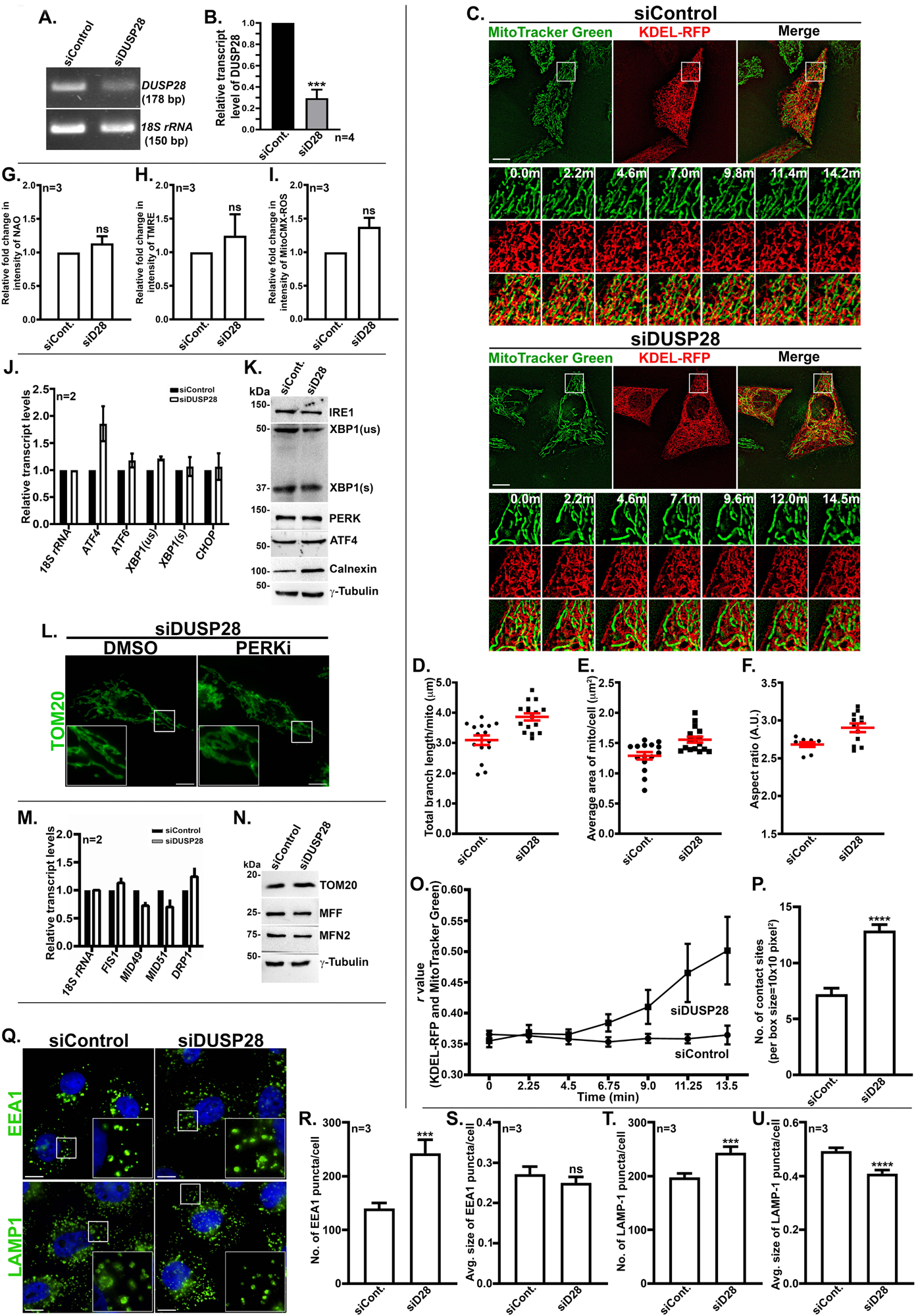
**(A)** Semiquantitative PCR analysis of DUSP28 levels in the siControl and siDUSP28 transfected HeLa cells. *18S rRNA* is used as the internal control. bp – base pairs. **(B)** Graph representing the relative transcript levels of the DUSP28 gene quantified from the agarose gel images. n=4. ***p≤0.001. **(C)** Time-lapse live cell imaging of siControl and siDUSP28 cells transfected with KDEL-RFP and stained with MitoTracker Green. Scale bar, 10 μm. The insets represent the white boxed area of the cell, shown at different time intervals as indicated. **(D-F, O-P)** Graphs represent the quantification of various parameters of live cell imaging shown in **C**: total branch length per mitochondria per cell **(D)**, average area of mitochondria per cell **(E)**, aspect ratio (arbitrary units, A.U.) of mitochondria per cell **(F)**, Pearson’s correlation coefficient (*r*-value) between KDEL-RFP and MitoTracker Green over 0 to 15 mins time frame **(O),** and number of contact sites between KDEL-RFP and MitoTracker Green per box size of 10*10 pixel^2^ **(P).** n=10-15 cells. **(G-I)** Graphs indicating the FACS analysis of cells treated with nonyl acridine orange (NAO; 100 nM 15 mins) **(G)**, Tetramethylrhodamine, ethyl ester (TMRE; 15 nM for 15 mins) **(H),** or MitoTracker Red CMX-ROS (MitoCMX-ROS; 100 nM 15 mins) **(I)** dyes. The relative fluorescence intensities were measured and plotted to illustrate the status of mitochondrial mass and potential in siDUSP28 compared to siControl cells. n=3. **(J)** Graph indicates the relative transcript levels of UPR downstream genes. *18S rRNA* is used as a control. n=2. **(K)** Immunoblotting analysis of cell lysates from siControl and siDUSP28 cells for the expression of ER and UPR specific proteins. γ-tubulin is used as a loading control**. (L)** IFM images of siDUSP28 cells treated with DMSO or PERKi, fixed and stained for mitochondria with anti-TOM20 antibody. **(M)** Graph representing the relative transcript values of mitochondrial genes in siControl and siDUSP28 cells obtained by semiquantitative RT-PCR. n=2. **(N)** Immunoblotting analysis of mitochondrial proteins in siControl and siDUSP28 cells. γ-tubulin is used as a loading control. **(Q)** IFM images of siControl and siDUSP28 cells stained for the markers of early endosomes (EEA1) and lysosomes (LAMP1). Insets of IFM images are amplified white box areas. Scale bar, 10 μm. **(R-U)** Graphs represent the quantification of the size and number of organelles in siControl and siDUSP28 cells. The number of EEA1 puncta per cell **®,** average size of EEA1 puncta per cell **(S),** number of LAMP1 puncta per cell **(T)** and the average size of LAMP1 puncta per cell **(U)** were measured in the cells shown in **Q**. n=3. All values are presented in the graphs as mean±s.e.m. ***p≤0.001, ****p≤0.0001 and, ns, non-significant.

**Figure S4.**
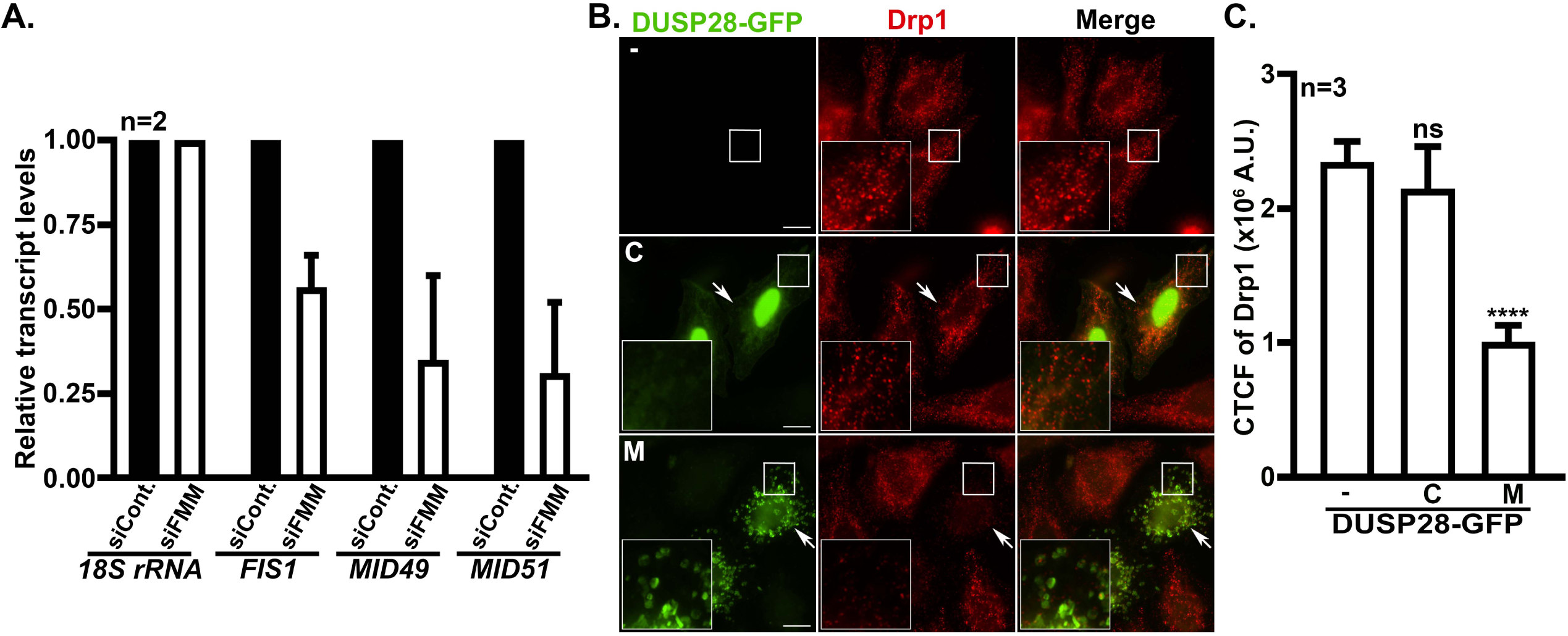
**(A)** Graph indicates the relative transcript values of *FIS1*, *MID49* and *MID51* genes in triple siRNA (siFIS1+siMID49+siMID51, siFMM) transfected HeLa cells compared to siControl cells. Transcripts were measured by RT-PCR, and *18S rRNA* is used as the control. n=2. **(B)** IFM images of untransfected and transfected HeLa cells with DUSP28-GFP, fixed and stained for Drp1. Images represent the fluorescence intensity of Drp1 in plain cells or cytosolic (C) or mitochondrial (M) localized DUSP28-GFP cells. Arrows point to the reduced Drp1 intensity in mitochondrial localized DUSP28-GFP cells. Insets are amplified white box areas. Scale bar, 10 μm. **(C)** Graph indicates the quantification of Drp1 fluorescence intensity (measured by corrected total cell fluorescence, A.U. – arbitrary units) in the cells shown in **B**. n=3. All values are presented in the graphs as mean±s.e.m. ****p≤0.0001 and ns, non-significant.

**Figure S5.**
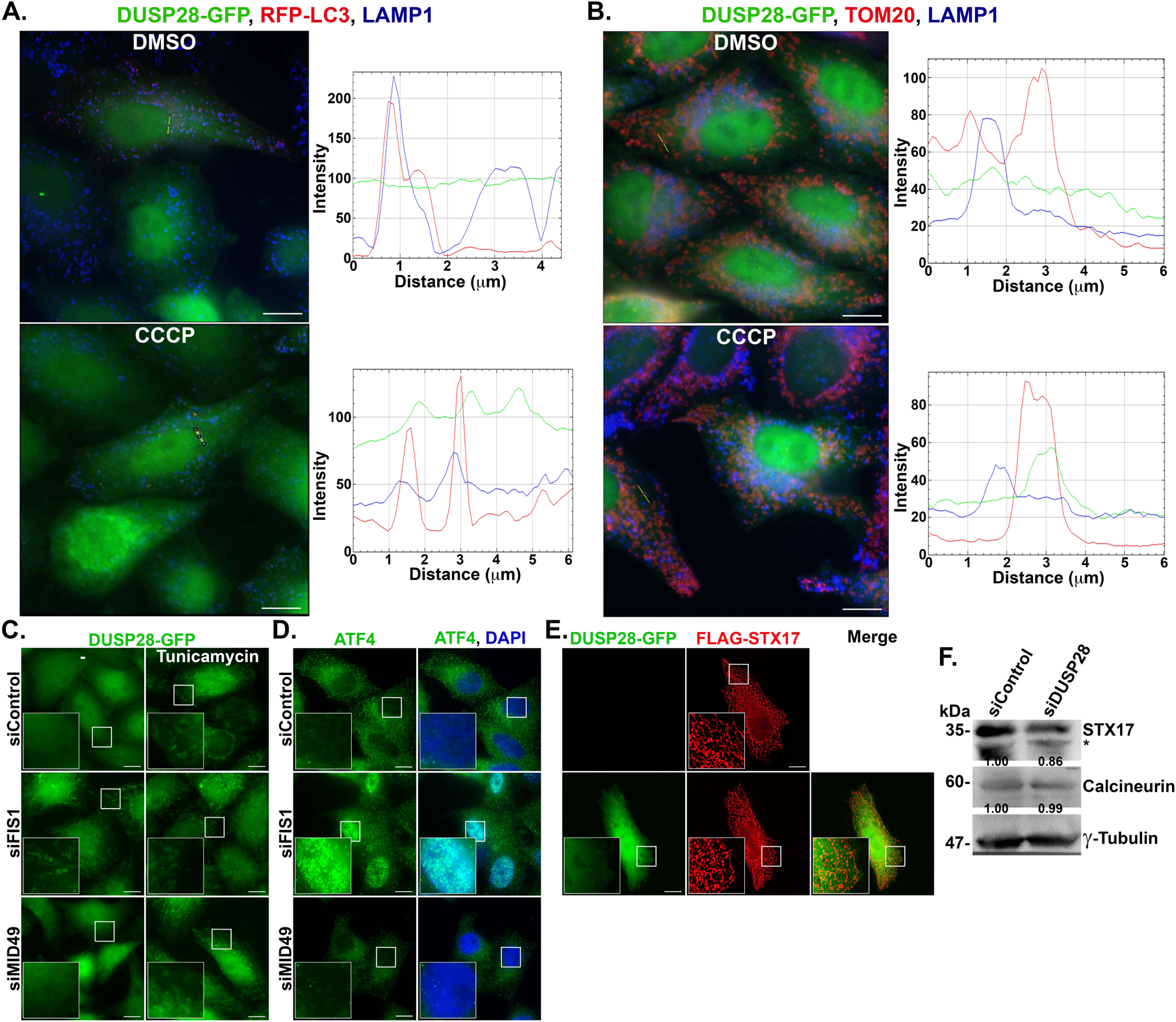
**(A)** IFM images of stable DUSP28-GFP expressing HeLa cells, transfected with RFP-LC3 followed by fixation and staining with anti-LAMP1 antibody (blue). (**B**) IFM images of stable DUSP28-GFP expressing HeLa cells fixed and stained with anti-TOM20 (red) and anti-LAMP1 antibodies (blue). In **A** and **B**, cells were treated with DMSO or CCCP. Scale bar, 10 μm. The graphs represent the intensity line plot for the proteins DUSP28-GFP, LC3-RFP/TOM20 and LAMP1 (green, red and blue lines correspondingly) in DMSO and CCCP treated cells. Note the merging of all three colours at certain points of the line in the CCCP-treated compared to the DMSO condition. **(C)** IFM images of stable HeLa cells expressing DUSP28-GFP, transfected with siControl, siFIS1 or siMID49 siRNA. Cells were treated with DMSO or tunicamycin to examine the localization of DUSP28-GFP. **(D)** IFM images of HeLa cells transfected with siControl, siFIS1 or siMID49 siRNA. Cells were fixed and stained with anti-ATF4 antibody (green) and DAPI (blue) to examine the nuclear localization of ATF4. **(E)** IFM images of HeLa cells transfected with FLAG-STX17 alone or combined with DUSP28-GFP to demonstrate the localization of STX17 in cells expressing cytosolic DUSP28-GFP. Insets of IFM images are amplified white box areas. Scale bar, 10 μm. **(F)** Immunoblotting of siControl and siDUSP28 HeLa cell lysates for the expression of STX17 and calcineurin. γ-tubulin is used as the loading control. *-represents the non-specific band developed with anti-STX17 antibody. Relative protein levels were indicated on the blots.

## Supplementary Videos

**Video S1**. The live-cell imaging of HeLa cells expressing DUSP28-GFP and mitochondrial marker DsRed2-Mito7. Cells were imaged for 4-5 min.

**Video S2**. The live-cell imaging of siControl treated HeLa cells expressing KDEL-RFP and stained with mitoTracker Green. Cells were imaged for 4-5 min.

**Video S3**. The live-cell imaging of siDUSP28 treated HeLa cells expressing KDEL-RFP and stained with mitoTracker Green. Cells were imaged for 4-5 min.

## References

Abrisch, R.G., S.C. Gumbin, B.T. Wisniewski, L.L. Lackner, and G.K. Voeltz. 2020. Fission and fusion machineries converge at ER contact sites to regulate mitochondrial morphology. Journal of Cell Biology. 219.

Ali, M.M., T. Bagratuni, E.L. Davenport, P.R. Nowak, M.C. Silva-Santisteban, A. Hardcastle, C. McAndrews, M.G. Rowlands, G.J. Morgan, W. Aherne, I. Collins, F.E. Davies, and L.H. Pearl. 2011. Structure of the Ire1 autophosphorylation complex and implications for the unfolded protein response. EMBO J. 30:894–905.

Almeida, L.M., B.R. Pinho, M.R. Duchen, and J.M.A. Oliveira. 2022. The PERKs of mitochondria protection during stress: insights for PERK modulation in neurodegenerative and metabolic diseases. Biol Rev Camb Philos Soc. 97:1737–1748.

Alonso, A., and R. Pulido. 2016. The extended human PTPome: a growing tyrosine phosphatase family. FEBS J. 283:2197–2201.

Alonso, A., J. Sasin, N. Bottini, I. Friedberg, I. Friedberg, A. Osterman, A. Godzik, T. Hunter, J. Dixon, and T. Mustelin. 2004. Protein tyrosine phosphatases in the human genome. Cell. 117:699–711.

Ashrafi, G., and T.L. Schwarz. 2013. The pathways of mitophagy for quality control and clearance of mitochondria. Cell Death Differ. 20:31–42.

Bakunts, A., A. Orsi, M. Vitale, A. Cattaneo, F. Lari, L. Tade, R. Sitia, A. Raimondi, A. Bachi, and E. van Anken. 2017. Ratiometric sensing of BiP-client versus BiP levels by the unfolded protein response determines its signaling amplitude. Elife. 6.

Borgese, N., M. Francolini, and E. Snapp. 2006. Endoplasmic reticulum architecture: structures in flux. Curr Opin Cell Biol. 18:358–364.

Brand, C.S., V.P. Tan, J.H. Brown, and S. Miyamoto. 2018. RhoA regulates Drp1 mediated mitochondrial fission through ROCK to protect cardiomyocytes. Cell Signal. 50:48–57.

Buiga, P., A. Elson, L. Tabernero, and J.M. Schwartz. 2018. Regulation of dual specificity phosphatases in breast cancer during initial treatment with Herceptin: a Boolean model analysis. BMC Syst Biol. 12:11.

Cereghetti, G.M., A. Stangherlin, O. Martins de Brito, C.R. Chang, C. Blackstone, P. Bernardi, and L. Scorrano. 2008. Dephosphorylation by calcineurin regulates translocation of Drp1 to mitochondria. Proc Natl Acad Sci U S A. 105:15803–15808.

Chang, C.R., and C. Blackstone. 2007. Cyclic AMP-dependent protein kinase phosphorylation of Drp1 regulates its GTPase activity and mitochondrial morphology. J Biol Chem. 282:21583–21587.

Chaudhry, A., R. Shi, and D.S. Luciani. 2020. A pipeline for multidimensional confocal analysis of mitochondrial morphology, function, and dynamics in pancreatic beta-cells. Am J Physiol Endocrinol Metab. 318:E87–E101.

Chen, Y., and F. Brandizzi. 2013. IRE1: ER stress sensor and cell fate executor. Trends Cell Biol. 23:547–555.

Cribbs, J.T., and S. Strack. 2007. Reversible phosphorylation of Drp1 by cyclic AMP-dependent protein kinase and calcineurin regulates mitochondrial fission and cell death. EMBO Rep. 8:939–944.

Csordas, G., D. Weaver, and G. Hajnoczky. 2018. Endoplasmic Reticulum-Mitochondrial Contactology: Structure and Signaling Functions. Trends Cell Biol. 28:523–540.

Dumont, F.J. 2000. FK506, an immunosuppressant targeting calcineurin function. Curr Med Chem. 7:731–748.

Elgass, K., J. Pakay, M.T. Ryan, and C.S. Palmer. 2013. Recent advances into the understanding of mitochondrial fission. Biochim Biophys Acta. 1833:150–161.

Forrest, A.R., T. Ravasi, D. Taylor, T. Huber, D.A. Hume, S. Grimmond, R.G. Group, and G.S.L. Members. 2003. Phosphoregulators: protein kinases and protein phosphatases of mouse. Genome Res. 13:1443–1454.

Friedman, J.R., L.L. Lackner, M. West, J.R. DiBenedetto, J. Nunnari, and G.K. Voeltz. 2011. ER tubules mark sites of mitochondrial division. Science. 334:358–362.

Friedman, J.R., and G.K. Voeltz. 2011. The ER in 3D: a multifunctional dynamic membrane network. Trends Cell Biol. 21:709–717.

Frohlich, C., S. Grabiger, D. Schwefel, K. Faelber, E. Rosenbaum, J. Mears, O. Rocks, and O. Daumke. 2013. Structural insights into oligomerization and mitochondrial remodelling of dynamin 1-like protein. EMBO J. 32:1280–1292.

Gallagher, C.M., C. Garri, E.L. Cain, K.K. Ang, C.G. Wilson, S. Chen, B.R. Hearn, P. Jaishankar, A. Aranda-Diaz, M.R. Arkin, A.R. Renslo, and P. Walter. 2016. Ceapins are a new class of unfolded protein response inhibitors, selectively targeting the ATF6alpha branch. Elife. 5.

Giacomello, M., A. Pyakurel, C. Glytsou, and L. Scorrano. 2020. The cell biology of mitochondrial membrane dynamics. Nat Rev Mol Cell Biol. 21:204–224.

Han, H., J. Tan, R. Wang, H. Wan, Y. He, X. Yan, J. Guo, Q. Gao, J. Li, S. Shang, F. Chen, R. Tian, W. Liu, L. Liao, B. Tang, and Z. Zhang. 2020. PINK1 phosphorylates Drp1(S616) to regulate mitophagy-independent mitochondrial dynamics. EMBO Rep. 21:e48686.

Han, X.J., Y.F. Lu, S.A. Li, T. Kaitsuka, Y. Sato, K. Tomizawa, A.C. Nairn, K. Takei, H. Matsui, and M. Matsushita. 2008. CaM kinase I alpha-induced phosphorylation of Drp1 regulates mitochondrial morphology. J Cell Biol. 182:573–585.

Hetz, C., and L.H. Glimcher. 2009. Fine-tuning of the unfolded protein response: Assembling the IRE1alpha interactome. Mol Cell. 35:551–561.

Ji, W.K., A.L. Hatch, R.A. Merrill, S. Strack, and H.N. Higgs. 2015. Actin filaments target the oligomeric maturation of the dynamin GTPase Drp1 to mitochondrial fission sites. Elife. 4:e11553.

Kar, U.P., H. Dey, and A. Rahaman. 2017. Regulation of dynamin family proteins by post-translational modifications. J Biosci. 42:333–344.

Kim, S.J., G.H. Syed, and A. Siddiqui. 2013. Hepatitis C virus induces the mitochondrial translocation of Parkin and subsequent mitophagy. PLoS Pathog. 9:e1003285.

Ko, A.R., H.W. Hyun, S.J. Min, and J.E. Kim. 2016. The Differential DRP1 Phosphorylation and Mitochondrial Dynamics in the Regional Specific Astroglial Death Induced by Status Epilepticus. Front Cell Neurosci. 10:124.

Kondapalli, C., A. Kazlauskaite, N. Zhang, H.I. Woodroof, D.G. Campbell, R. Gourlay, L. Burchell, H. Walden, T.J. Macartney, M. Deak, A. Knebel, D.R. Alessi, and M.M. Muqit. 2012. PINK1 is activated by mitochondrial membrane potential depolarization and stimulates Parkin E3 ligase activity by phosphorylating Serine 65. Open Biol. 2:120080.

Ku, B., W. Hong, C.W. Keum, M. Kim, H. Ryu, D. Jeon, H.C. Shin, J.H. Kim, S.J. Kim, and S.E. Ryu. 2017. Structural and biochemical analysis of atypically low dephosphorylating activity of human dual-specificity phosphatase 28. PLoS One. 12:e0187701.

Lebeau, J., J.M. Saunders, V.W.R. Moraes, A. Madhavan, N. Madrazo, M.C. Anthony, and R.L. Wiseman. 2018. The PERK Arm of the Unfolded Protein Response Regulates Mitochondrial Morphology during Acute Endoplasmic Reticulum Stress. Cell Rep. 22:2827–2836.

Lee, J., J. Hun Yun, J. Lee, C. Choi, and J. Hoon Kim. 2015. Blockade of dual-specificity phosphatase 28 decreases chemo-resistance and migration in human pancreatic cancer cells. Sci Rep. 5:12296.

Lee, J., J. Lee, and J.H. Kim. 2019. Scattered DUSP28 is a novel biomarker responsible for aggravating malignancy via the autocrine and paracrine signaling in metastatic pancreatic cancer. Cancer Lett. 456:1–12.

Lee, J., J. Lee, J.H. Yun, C. Choi, S. Cho, S.J. Kim, and J.H. Kim. 2017. Autocrine DUSP28 signaling mediates pancreatic cancer malignancy via regulation of PDGF-A. Sci Rep. 7:12760.

Lin, J.H., H. Li, Y. Zhang, D. Ron, and P. Walter. 2009. Divergent effects of PERK and IRE1 signaling on cell viability. PLoS One. 4:e4170.

Loson, O.C., Z. Song, H. Chen, and D.C. Chan. 2013. Fis1, Mff, MiD49, and MiD51 mediate Drp1 recruitment in mitochondrial fission. Mol Biol Cell. 24:659–667.

Mahameed, M., T. Wilhelm, O. Darawshi, A. Obiedat, W.S. Tommy, C. Chintha, T. Schubert, A. Samali, E. Chevet, L.A. Eriksson, M. Huber, and B. Tirosh. 2019. The unfolded protein response modulators GSK2606414 and KIRA6 are potent KIT inhibitors. Cell Death Dis. 10:300.

Malhotra, J.D., and R.J. Kaufman. 2007. Endoplasmic reticulum stress and oxidative stress: a vicious cycle or a double-edged sword? Antioxid Redox Signal. 9:2277–2293.

Manning, G., D.B. Whyte, R. Martinez, T. Hunter, and S. Sudarsanam. 2002. The protein kinase complement of the human genome. Science. 298:1912–1934.

McQuiston, A., and J.A. Diehl. 2017. Recent insights into PERK-dependent signaling from the stressed endoplasmic reticulum. F1000Res. 6:1897.

Mori, K. 2009. Signalling pathways in the unfolded protein response: development from yeast to mammals. J Biochem. 146:743–750.

Munoz, J.P., S. Ivanova, J. Sanchez-Wandelmer, P. Martinez-Cristobal, E. Noguera, A. Sancho, A. Diaz-Ramos, M.I. Hernandez-Alvarez, D. Sebastian, C. Mauvezin, M. Palacin, and A. Zorzano. 2013. Mfn2 modulates the UPR and mitochondrial function via repression of PERK. EMBO J. 32:2348–2361.

Nag, S., K. Szederkenyi, O. Gorbenko, H. Tyrrell, C.M. Yip, and G.A. McQuibban. 2023. PGAM5 is an MFN2 phosphatase that plays an essential role in the regulation of mitochondrial dynamics. Cell Rep. 42:112895.

Narendra, D.P., S.M. Jin, A. Tanaka, D.F. Suen, C.A. Gautier, J. Shen, M.R. Cookson, and R.J. Youle. 2010. PINK1 is selectively stabilized on impaired mitochondria to activate Parkin. PLoS Biol. 8:e1000298.

Otera, H., and K. Mihara. 2011. Discovery of the membrane receptor for mitochondrial fission GTPase Drp1. Small GTPases. 2:167–172.

Padman, B.S., M. Bach, G. Lucarelli, M. Prescott, and G. Ramm. 2013. The protonophore CCCP interferes with lysosomal degradation of autophagic cargo in yeast and mammalian cells. Autophagy. 9:1862–1875.

Palmer, C.S., L.D. Osellame, D. Laine, O.S. Koutsopoulos, A.E. Frazier, and M.T. Ryan. 2011. MiD49 and MiD51, new components of the mitochondrial fission machinery. EMBO Rep. 12:565–573.

Patterson, K.I., T. Brummer, P.M. O’Brien, and R.J. Daly. 2009. Dual-specificity phosphatases: critical regulators with diverse cellular targets. Biochem J. 418:475–489.

Peng, J., T. Wang, C. Yue, X. Luo, and P. Xiao. 2022. PGAM5: A necroptosis gene associated with poor tumor prognosis that promotes cutaneous melanoma progression. Front Oncol. 12:1004511.

Perea, V., C. Cole, J. Lebeau, V. Dolina, K.R. Baron, A. Madhavan, J.W. Kelly, D.A. Grotjahn, and R.L. Wiseman. 2023. PERK signaling promotes mitochondrial elongation by remodeling membrane phosphatidic acid. EMBO J. 42:e113908.

Pulido, R., and R. Hooft van Huijsduijnen. 2008. Protein tyrosine phosphatases: dual-specificity phosphatases in health and disease. FEBS J. 275:848–866.

Pulido, R., and R. Lang. 2019. Dual Specificity Phosphatases: From Molecular Mechanisms to Biological Function. Int J Mol Sci. 20.

Rowland, A.A., and G.K. Voeltz. 2012. Endoplasmic reticulum-mitochondria contacts: function of the junction. Nat Rev Mol Cell Biol. 13:607–625.

Sacco, F., L. Perfetto, L. Castagnoli, and G. Cesareni. 2012. The human phosphatase interactome: An intricate family portrait. FEBS Lett. 586:2732–2739.

Schuck, S., W.A. Prinz, K.S. Thorn, C. Voss, and P. Walter. 2009. Membrane expansion alleviates endoplasmic reticulum stress independently of the unfolded protein response. J Cell Biol. 187:525–536.

Schwarz, D.S., and M.D. Blower. 2016. The endoplasmic reticulum: structure, function and response to cellular signaling. Cell Mol Life Sci. 73:79–94.

Shakya, S., P. Sharma, A.M. Bhatt, R.A. Jani, C. Delevoye, and S.R. Setty. 2018. Rab22A recruits BLOC-1 and BLOC-2 to promote the biogenesis of recycling endosomes. EMBO Rep. 19.

Shibata, Y., G.K. Voeltz, and T.A. Rapoport. 2006. Rough sheets and smooth tubules. Cell. 126:435–439.

Singh, V., C. Erady, and N. Balasubramanian. 2018. Cell-matrix adhesion controls Golgi organization and function through Arf1 activation in anchorage-dependent cells. J Cell Sci. 131.

Smirnova, E., L. Griparic, D.L. Shurland, and A.M. van der Bliek. 2001. Dynamin-related protein Drp1 is required for mitochondrial division in mammalian cells. Mol Biol Cell. 12:2245–2256.

Spinelli, J.B., and M.C. Haigis. 2018. The multifaceted contributions of mitochondria to cellular metabolism. Nat Cell Biol. 20:745–754.

Sugo, M., H. Kimura, K. Arasaki, T. Amemiya, N. Hirota, N. Dohmae, Y. Imai, T. Inoshita, K. Shiba-Fukushima, N. Hattori, J. Cheng, T. Fujimoto, Y. Wakana, H. Inoue, and M. Tagaya. 2018. Syntaxin 17 regulates the localization and function of PGAM5 in mitochondrial division and mitophagy. EMBO J. 37.

Tonks, N.K. 2006. Protein tyrosine phosphatases: from genes, to function, to disease. Nat Rev Mol Cell Biol. 7:833–846.

van der Bliek, A.M., Q. Shen, and S. Kawajiri. 2013. Mechanisms of mitochondrial fission and fusion. Cold Spring Harb Perspect Biol. 5.

Verfaillie, T., N. Rubio, A.D. Garg, G. Bultynck, R. Rizzuto, J.P. Decuypere, J. Piette, C. Linehan, S. Gupta, A. Samali, and P. Agostinis. 2012. PERK is required at the ER-mitochondrial contact sites to convey apoptosis after ROS-based ER stress. Cell Death Differ. 19:1880–1891.

Vives-Bauza, C., C. Zhou, Y. Huang, M. Cui, R.L. de Vries, J. Kim, J. May, M.A. Tocilescu, W. Liu, H.S. Ko, J. Magrane, D.J. Moore, V.L. Dawson, R. Grailhe, T.M. Dawson, C. Li, K. Tieu, and S. Przedborski. 2010. PINK1-dependent recruitment of Parkin to mitochondria in mitophagy. Proc Natl Acad Sci U S A. 107:378–383.

Walter, P., and D. Ron. 2011. The unfolded protein response: from stress pathway to homeostatic regulation. Science. 334:1081–1086.

Wang, D., S. Han, R. Peng, C. Jiao, X. Wang, Z. Han, and X. Li. 2014. DUSP28 contributes to human hepatocellular carcinoma via regulation of the p38 MAPK signaling. Int J Oncol. 45:2596–2604.

Wang, Z., H. Jiang, S. Chen, F. Du, and X. Wang. 2012. The mitochondrial phosphatase PGAM5 functions at the convergence point of multiple necrotic death pathways. Cell. 148:228–243.

Wieckowski, M.R., C. Giorgi, M. Lebiedzinska, J. Duszynski, and P. Pinton. 2009. Isolation of mitochondria-associated membranes and mitochondria from animal tissues and cells. Nat Protoc. 4:1582–1590.

Xian, H., Q. Yang, L. Xiao, H.M. Shen, and Y.C. Liou. 2019. STX17 dynamically regulated by Fis1 induces mitophagy via hierarchical macroautophagic mechanism. Nat Commun. 10:2059.

Yoon, Y., E.W. Krueger, B.J. Oswald, and M.A. McNiven. 2003. The mitochondrial protein hFis1 regulates mitochondrial fission in mammalian cells through an interaction with the dynamin-like protein DLP1. Mol Cell Biol. 23:5409–5420.

Yu, R., S.B. Jin, M. Ankarcrona, U. Lendahl, M. Nister, and J. Zhao. 2021. The Molecular Assembly State of Drp1 Controls its Association With the Mitochondrial Recruitment Receptors Mff and MIEF1/2. Front Cell Dev Biol. 9:706687.

Zaja, I., X. Bai, Y. Liu, C. Kikuchi, S. Dosenovic, Y. Yan, S.G. Canfield, and Z.J. Bosnjak. 2014. Cdk1, PKCdelta and calcineurin-mediated Drp1 pathway contributes to mitochondrial fission-induced cardiomyocyte death. Biochem Biophys Res Commun. 453:710–721.

